# The Alzheimer’s-Associated *SORL1* p.Y1816C Variant Impairs APP Sorting, Axonal Trafficking, and Neuronal Activity in iPSC-Derived Brain Models

**DOI:** 10.1101/2025.07.28.667160

**Authors:** Jan Raska, Miriam Satkova, Klara Plesingrova, Jiri Sedmik, Sona Cesnarikova, Monica Feole, Ondrej Bernatik, Victorio M. Pozo Devoto, Hana Hribkova, Katerina Amruz Cerna, Nadezda Vaskovicova, Simona Bartova, Veronika Pospisilova, Tereza Vanova, Petr Fojtik, Olav M. Andersen, Dasa Bohaciakova

**Affiliations:** Masaryk University, Faculty of Medicine, Department of Histology and Embryology, Brno, Czech Republic; International Clinical Research Center (ICRC), St. Anne’s University Hospital, Brno, Czech Republic; Aarhus University, Department of Biomedicine, Aarhus, Denmark

**Keywords:** SORL1, SORLA, Alzheimer’s disease, Endosomal trafficking, APP processing, Axonal transport and live-cell imaging, Neuronal electrophysiology and hyperexcitability, iPSC-derived neurons, Neural organoids, Aβ peptides

## Abstract

**Background:** *SORL1*, encoding the sorting receptor SORLA, is now recognized as the fourth autosomal dominant Alzheimer’s disease (AD) gene. Loss of SORLA function is known to disrupt endosomal trafficking and enhance amyloidogenic APP processing, two key aspects of the onset and progression of AD. However, the pathogenic consequences of disrupted endolysosomal pathways, deregulated protein sorting, as well as the effects of specific *SORL1* missense variants on human neuronal function, still remain understudied.

**Methods:** Our investigations were performed using two complementary human iPSC-derived models: 2D NGN2-induced neurons and 3D cerebral organoids established from isogenic wild-type (WT), *SORL1* p.Y1816C (KI) missense variant, and SORL1 knock-out (KO) cells. We analyzed SORLA maturation and ectodomain shedding, APP localization, and amyloid-β secretion. Endosomal morphology and neuritic swellings were assessed via electron microscopy, while axonal transport of APP and Rab5+ endosomes was evaluated through live-cell imaging. Neuronal network activity was measured using multielectrode array recordings.

**Results:** Our results demonstrate that the p.Y1816C variant leads to impaired SORLA maturation and reduced shedding, without affecting neuronal or organoid differentiation. Notably, we show an ultrastructure of endosomes, including their content, and demonstrate that both KO and KI models exhibit early endosome enlargement, increased APP retention in endosomes, elevated Aβ40/42 secretion, and amyloid-β deposition in 3D organoids. Importantly, we identified previously uncharacterized functional consequences of abolished SORLA activity, including axonal swellings and significantly impaired transport of Rab5+ endosomes and APP, characterized by deregulated velocities, directionality of transport, and increased stalling. Additionally, we discovered that both KO and p.Y1816C KI neurons exhibit abnormal electrophysiological activity, including increased spontaneous firing, burst frequency, and network synchrony.

**Conclusions:** Our study defines the mechanistic consequences of the *SORL1* p.Y1816C variant and demonstrates its pathogenicity in human neurons. Importantly, we also identify novel roles for SORLA in maintaining axonal transport homeostasis and regulating neuronal excitability, expanding its functional relevance beyond endosomal APP processing. These findings reinforce the central role of endosomal trafficking disruption in AD and support the use of isogenic human models for evaluating AD risk variants.

## 1. Background

Alzheimer’s disease (AD), the most common neurodegenerative disorder and leading cause of dementia, poses a major burden for aging populations. And while hallmark features such as amyloid-β (Aβ) plaques and tau tangles remain central to AD pathology, increasing evidence implicates early intracellular trafficking defects as key contributors to disease progression. In particular, endosomal dysfunction is being recognized as a fundamental driver, detectable even before the onset of extracellular Aβ deposition [1–6]. Among AD-linked genes, *SORL1* has emerged as the fourth autosomal dominant gene alongside *APP*, *PSEN1*, and *PSEN2* [7–9]. It encodes SORLA, a neuronal sorting receptor essential for the trafficking of multiple cargos, most notably amyloid precursor protein (APP). In healthy neurons, SORLA binds APP and directs it away from early endosomes, where β-secretase (BACE1)-mediated amyloidogenic processing occurs, thereby limiting the generation of toxic Aβ peptides. [7,10]. Loss of SORLA function due to *SORL1* mutations leads to missorting of APP into BACE1-containing endosomes, resulting in enhanced Aβ production and secretion. These trafficking defects are often accompanied by a marked enlargement of early endosomes, a morphological feature found not only in *SORL1*-deficient models but also in other familial AD mutations, supporting its relevance as an early and generalizable AD-related phenotype [11–16]. As much of this knowledge derives from *SORL1* knock-out (KO) studies or overexpression models, the functional consequences of individual missense *SORL1* variants found in patients remain poorly understood.

Notably, we have previously described that one such variant, p.Y1816C, segregates with AD in three unrelated families [17]. It lies within the fibronectin type III (3Fn) domain of SORLA, which we found to be essential for dimerization and interaction with the retromer complex that mediates the endosome-to-surface trafficking of APP [18,19]. Extending these findings, we further provided evidence that the p.Y1816C mutant impairs SORLA homodimerization in the endosome, leading to decreased trafficking to the cell surface and less soluble SORLA (sSORLA) ectodomain shedding. These trafficking defects of the mutant receptor, described in the experimental cellular models with overexpressed SORLA, can be rescued by the expression of the SORLA 3Fn-minireceptor [17,18]. However, the broader impact of the p.Y1816C mutation on the maturation of SORLA at the endogenous level, trafficking dynamics, and functional role in APP processing within human neuronal and 3D microphysiological systems remained unanswered.

Importantly, neurons, being highly polarized, rely on the efficient long-range transport of key cargoes, such as APP and Rab5-positive (Rab5+) early endosomes, via microtubule-based motor proteins [2,20,21]. Disruptions in these processes are consistently observed in AD models, where axonal swellings containing vesicular accumulations appear in patient brains, animal models, and iPSC-derived neurons with familial AD mutations [2,20,22,23]. Notably, our recent study has shown that neurons carrying the APP Swedish mutation exhibit impaired vesicle motility and increased stalling of APP, contributing to synaptic dysfunction [19]. Despite these findings, the direct involvement of SORLA in axonal transport or maintenance of axonal integrity has not been defined.

In parallel, growing evidence links AD pathogenesis to early neuronal hyperexcitability. iPSC-derived neurons with *APP* or *PSEN1* mutations display increased spontaneous activity, altered calcium dynamics, and network synchrony [14,15,24]. Similar abnormalities, such as epileptiform discharges and increased seizure susceptibility, are also seen in 5xFAD and APP/PS1 mouse models [25,26]. Interestingly, Zhou *et al.* previously demonstrated that the APP Swedish mutation in iPSC-derived neurons increases the number of functional synapses and strongly suggested that, at the physiological level, Aβ might be synaptogenic [27]. Although SORLA is well established as a regulator of APP trafficking and Aβ production, its potential influence on neuronal excitability and network activity remains unclear. Aside from limited evidence of increased firing in *SORL1*-deficient neurons [15], the broader electrophysiological consequences of SORLA dysfunction have yet to be investigated.

To address these gaps, we generated isogenic human iPSC-derived neuronal and cerebral organoid models carrying either the *SORL1* p.Y1816C variant heterozygously or a full (homozygous) *SORL1* knock-out. We then systematically examined their molecular, structural, and functional phenotypes to determine whether this missense mutation, as well as the complete absence of SORLA, reproduces the key cellular hallmarks of AD. Our analyses reveal that the pathogenic *SORL1* p.Y1816C variant leads to impaired protein maturation and shedding. SORLA dysfunction then leads to enlarged endosomes with APP retention, increased Aβ secretion, disrupted axonal transport, neuritic swellings, and hyperactive neuronal networks. Altogether, these results highlight the multifaceted role of SORLA in maintaining neuronal health and underscore the importance of trafficking pathways as early and actionable drivers of AD.

## 2. Methods

### 2.1 Cell culture and gene editing

A human iPSC line with a stably integrated, doxycycline-inducible NGN2 transgene (i3N iPSCs, RRID: CVCL_C7XJ) was used in this study. The i3N cell line was generously provided by Michael Ward [28]. Cells were maintained as a monolayer on Matrigel® hESC-qualified matrix (Corning) in mTeSR™1 medium (STEMCELL Technologies) supplemented with 1X Zellshield™ (Minerva Biolabs) and passaged using TrypLE™ Express reagent (Thermo Fisher Scientific). IPSCs were maintained at 37 °C in a 5% CO_2_ incubator.

Three clones of *SORL1* Knock-Out (KO) and one clone of heterozygous p.Y1816C Knock-In (KI) iPSCs used in this study were generated using a CRISPR/Cas9 ribonucleoprotein (RNP) complex containing Cas9-GFP (Sigma). For *SORL1*-KO, exon 6 of the *SORL1* gene in i3N iPSCs was targeted using a previously published guide RNA (gRNA)[16]. The p.Y1816C KI was introduced by targeting exon 41 with a custom-designed gRNA and a single-stranded oligodeoxynucleotide (ssODN) as a template for homology-directed repair, as published previously [29]. CRISPR/Cas9 RNP complexes were delivered using the Neon™ Transfection System (Thermo Fisher Scientific), followed by GFP-based single-cell sorting into 96-well plates to generate clonal lines. *SORL1*-KO clones were initially screened by Western blotting for the absence of SORLA protein by LR11 antibody (BD Transduction Laboratories) and confirmed by Sanger sequencing (Seqme). The *SORL1*-KI clone was initially screened by PCR-RFLP. PCR product with WT sequence was cleaved by BtsCI (New England Biolabs), while PCR product with the introduced p.Y1816C mutation was cleaved by AciI (New England Biolabs) and confirmed by Sanger sequencing (Seqme). To control for the potential effects of the genome editing process itself, two MOCK control clones were generated by subjecting the parental WT iPSC line to the complete CRISPR workflow, including transfection, sorting, and clonal expansion, without introducing any targeted modification. Unless otherwise stated, all experiments were performed using three clones of WT iPSCs (one parental cell line and two MOCK clones), three clones of KO iPSCs, and one clone of KI iPSCs.

A comprehensive list of all used cell lines and clones is provided in **Table S1**; a list of all experimental samples, including iPSC clones used for individual analyses and the number of biological replicates, is provided in **Table S2;** and sequences of gRNAs, ssODNs, and PCR primers are provided in **Table S3**.

### 2.2 Differentiation of 2D neurons and 3D cerebral organoids

2D Neuronal differentiation of i3N iPSCs was performed using a previously published protocol [28]. Briefly, undifferentiated iPSCs (Day 0) were plated as a single-cell monolayer on Matrigel®-coated dishes at a density of 0.7 × 10⁵ cells/cm² in Induction Medium (IM) supplemented with 2 μg/mL doxycycline (Sigma) and 10 μM Y-27632 ROCK inhibitor (Selleckchem). From Day 1 to Day 2, IM containing 2 μg/mL doxycycline was changed daily. On Day 3, cells were replated onto Poly-L-Ornithine-coated dishes (Sigma) using Accutase (Thermo Fisher Scientific) at a density of 0.8 × 10⁵ cells/cm² in Cortical Neuron Medium (CNM). Half of the CNM was replaced twice per week until sample collection. Samples were harvested on Days 20, 30, and 40. For the collection of conditioned media for ELISA and Western blot analyses, CNM was replaced with Essential 6 medium (Thermo Fisher Scientific), and cells were cultured for an additional 3 days prior to sampling. Media composition is provided in **Table S3.**

3D cerebral organoids were generated using a modified protocol from [30] as previously described by Vanova *et al.* [31]. Briefly, on Day 0, iPSC lines were dissociated using Accutase (Thermo Fisher Scientific) and seeded into non-adherent, V-bottom 96-well plates at a density of 2,000 cells per well in 150 µl of mTeSR™1 medium (STEMCELL Technologies) supplemented with 50 μM Y-27632 (Selleckchem) to initiate embryoid body (EB) formation. The next day, the medium was changed to fresh mTeSR without Y-27632 (100 µl per well). Once EBs reached a diameter of ≥400 μm, fresh Neural Induction Medium (NIM) was added daily for 6 days (typically from Day 3 to Day 8, 100 µl per well). After this induction phase, organoids were embedded in 7 μL of cold Geltrex™ (Thermo Fisher Scientific). Following polymerization, the Geltrex droplets containing organoids were gently detached and cultured without agitation on Petri dishes in Cerebral Organoid Differentiation Medium without vitamin A (CODM−A) for 7 days. Organoids were then transferred to Cerebral Organoid Differentiation Medium with vitamin A (CODM+A) and placed on an orbital shaker between Days 24 and 28. Half of the medium was replaced three times per week. Samples were collected on Days 9, 15, 30, and 60. For whole cell lysate and RNA isolation, 5 to 10 organoids were pooled per sample. For ELISA, individual organoids were cultured in 12-well plates with 0.8-1.5 mL of Essential 6 medium (Thermo Fisher Scientific) for 72 hours prior to the collection of conditioned media and organoids. Media compositions are provided in **Table S3.**

### 2.3 DNA Isolation and PCR analysis of ApoE genotype

Genomic DNA was isolated using the DNeasy Blood & Tissue Kit (Qiagen) according to the manufacturer’s instructions. PCR amplification was performed with Taq DNA polymerase (Top-Bio). Apolipoprotein E (*ApoE*) polymorphisms were analyzed using an allele-specific PCR method as previously described [32]. *GDF15p* was used as a loading control. PCR products were separated by electrophoresis on a 2% agarose gel containing GelRed® (Biotium) and visualized using the ChemiDoc™ Touch Imaging System (Bio-Rad). Primer sequences are listed in **Table S3.**

### 2.4 RNA isolation, cDNA synthesis, and qPCR

Total RNA was extracted using RNA Blue reagent (Top-Bio) following the manufacturer’s protocol. RNA concentration and purity were assessed using a NanoDrop 1000 spectrophotometer (Thermo Fisher Scientific). Reverse transcription was performed with 0.5–1 μg of total RNA using the Transcriptor First Strand cDNA Synthesis Kit (Roche), according to the manufacturer’s instructions. Quantitative PCR (qPCR) was carried out using the LightCycler® 480 SYBR Green I Master Kit (Roche) in 10 μL reactions on a LightCycler® 480 II system (Roche). The thermal cycling protocol included an initial denaturation at 95 °C for 5 min, followed by 45 amplification cycles (95 °C for 10 s, 60 °C for 10 s, and 72 °C for 10 s). Threshold cycle (Ct) values were determined using the Second Derivative Maximum Method in the LightCycler® 480 software. Gene expression was analyzed using the ΔCt method (Ct_target – Ct_housekeeping) and reported as 2^–ΔCt. Primer sequences are listed in **Table S3**.

### 2.5 Western blotting

Cells were washed with ice-cold PBS and lysed on ice for 1 hour in lysis buffer (20 mM Tris-HCl, pH 8.1; 10 mM EDTA; 1% Triton X-100; 1% NP-40) supplemented with cOmplete™ Mini Protease Inhibitor (Roche). Samples were sonicated by SonoPuls HD 2200 (power 20%; pulse 1s; time 10s; Bandelin). Protein concentrations were determined using the DC™ Protein Assay Kit (Bio-Rad). Equal amounts of protein were mixed with 4× Laemmli buffer, heated at 95 °C for 10 minutes, and separated by SDS-PAGE using custom-cast acrylamide gels. Proteins were transferred to PVDF membranes (Merck). Membranes were blocked for 1 hour in blocking buffer (5% skim milk in TBS containing 0.1% Tween-20), followed by overnight incubation at 4°C with primary antibodies diluted in blocking buffer. The following day, membranes were incubated with HRP-conjugated secondary antibodies for 1 hour at room temperature. Signals were detected using Clarity™ and Clarity Max™ ECL substrates (Bio-Rad) and visualized with the ChemiDoc™ Touch Imaging System (Bio-Rad). Band intensities were quantified using Image Lab software (Bio-Rad). A complete list of antibodies is provided in **Table S3.**

### 2.6 ELISA

Conditioned cell culture media samples were centrifuged at 10,000 × g for 5 minutes, and the resulting supernatant was collected, supplemented with cOmplete™ Mini Protease Inhibitor (Roche), and stored at –80 °C until ELISA analysis. Media volumes (V) were measured at the time of collection to account for the variable rate of evaporation between wells, and the values were used for ELISA normalization. Whole-cell lysates were prepared by incubating cells on ice for 1 hour in lysis buffer (20mM Tris-HCl, pH 8.1; 10m EDTA; 1% Triton X-100; 1% NP-40) supplemented with cOmplete™ Mini Protease Inhibitor (Roche). Total protein concentration in the lysates was determined using the DC Protein Assay Kit (Bio-Rad), multiplied by lysate volume to obtain total protein mass (m), and used to normalize ELISA results. Levels of Aβ peptides (Aβ40 and Aβ42) in conditioned media were quantified using the Amyloid Beta 40 Human ELISA Kit and the Amyloid Beta 42 Human ELISA Kit, Ultrasensitive (both Thermo Fisher Scientific), according to the manufacturer’s protocols. SORLA levels in conditioned media were measured using the Human SORL1 ELISA Kit (Abcam). All samples were analyzed in technical duplicates. To enable comparison across samples, values obtained from ELISA (c) were normalized to total protein content and media volume using the formula (c*V)/m and are expressed as picograms (pg) of analyte per microgram (µg) of total protein.

### 2.7 Immunocytochemistry and image analysis

iPSC-derived 2D neurons were seeded at a density of 75,000 cells per well on glass coverslips (Thermo Fisher Scientific) placed in 24-well plates. Cells were fixed in 3.7% paraformaldehyde for 20 minutes and permeabilized with 0.1% Triton X-100, followed by blocking in a solution containing 1% BSA and 0.03% Tween-20 in PBS. Primary antibodies diluted in blocking buffer were applied overnight at 4 °C. The next day, cells were incubated with secondary antibodies for 1 hour at room temperature. A complete list of antibodies is provided in **Table S3**. Nuclei were stained with DAPI (Sigma), and coverslips were mounted using ProLong™ Gold Antifade (Thermo Fisher Scientific). Imaging was performed using a Zeiss LSM 800 laser scanning confocal microscope equipped with a 561 nm laser and a Plan-Apochromat 63×/1.40 oil immersion objective (Zeiss). Image analysis was conducted using IMARIS software v.10 (Oxford Instruments). The “Surfaces” tool was used to evaluate endosome size, with volume selected as the measurement parameter. Thresholding was manually calibrated using control samples, and the same parameters were applied for batch analysis of all datasets. For data analysis, a total of 12 images per genotype were examined. This included four biological replicates per genotype, with three images analyzed per replicate.

For 3D organoid analyses, harvested samples were fixed with 3.7% paraformaldehyde for 1 hour at room temperature. Fixed COs were embedded in 4% agarose and sectioned into 200 μm slices using vibratome (Leica). Slices were incubated for 4 hours at room temperature in a blocking buffer containing 0.5% Triton X-100, 5% BSA, and 0.01% sodium azide in PBS. Primary antibodies, diluted in blocking buffer, were applied for 4 days at 4°C, followed by incubation with secondary antibodies for 24 hours at 4 °C. A complete list of antibodies is provided in **Table S3**. Nuclei were stained with DAPI (Sigma), and sections were cleared using a glycerol/fructose solution to enhance imaging depth and quality. Microscopy was performed using Axio Observer.Z1 microscope equipped with an LSM800 confocal unit and a Plan-Apochromat 20×/0.8 air objective (Zeiss). The pinhole size was set to 1 Airy unit. Fluorescent signals from labelled proteins and nuclei were acquired using high-resolution confocal settings and appropriate laser/filter combinations. Image analysis was performed using IMARIS software version 10 (Oxford Instruments). The “Surfaces” tool was used to assess APP size, with volume selected as the measurement parameter. Thresholding was manually calibrated using control samples, and the same parameters were applied consistently for batch analysis of all datasets. For data analysis, a total of 108 images per genotype were examined. This included three biological replicates per genotype, with each replicate comprising three organoids, and three sections analyzed from each organoid.

### 2.8 Proximity Ligation Assay

iPSC-derived neurons were seeded at a density of 75,000 cells per well on glass coverslips (Thermo Fisher Scientific) placed in 24-well plates. Cells were fixed and permeabilized as described for Immunofluorescence and processed for PLA using the Duolink® In Situ Red Starter Kit Mouse/Rabbit (Sigma) according to the manufacturer’s instructions. Cells were incubated in the provided blocking buffer and subsequently with primary antibodies overnight at 4 °C, using concentrations listed in **Table S3**. Following incubation with PLA probes, the ligation and amplification steps were carried out according to the kit protocol. Cells were then mounted in Duolink® In Situ Mounting Medium with DAPI. Images were acquired within two days of completing the PLA procedure to ensure signal preservation. Microscopy was performed using Axio Observer.Z1 microscope equipped with an LSM800 confocal unit and a Plan-Apochromat 63×/1.40 oil immersion objective (Zeiss). Images were captured at a resolution of 1024 × 1024 pixels. Image analysis was performed using ImageJ software by the Analyze Particles tool (size=10-150 pixel; circularity=0.00-1.00; optimized for 63X). For data analysis, a total of 39 images per genotype were examined. This included three biological replicates per genotype, with 13 images analyzed per replicate.

### 2.9 Transmission electron microscopy – TEM

iPSC-derived neurons were seeded at a density of 300,000 cells per well on a 6-well plate. After 20 days of maturation in CNM medium, samples were fixed in 3% glutaraldehyde (Sigma) in 0.1 M cacodylate buffer (pH 7.3; Sigma) with 2% saccharose and 1% tannin for 24 h at 4°C. After washing by 0.1M cacodylate buffer samples were post-fixed in 1% OsO4 (Sigma) in the same buffer for 1 h at room temperature, following by dehydration ethanol series (50%, 70%, 96% and 100% ethanol) and embedded in the LR White resin (Agar Scientific), polymerization for 3 days at 65°C. The resin blocks were processed using the standard protocol for electron microscopy, with cutting performed on a LEICA EM UC6 ultramicrotome (Leica). The ultrathin sections were examined using Morgagni 268D transmission electron microscope (Thermo Fisher Scientific), working at 90 kV and equipped with Veleta CCD camera (Olympus). For data analysis, a total of 60 images per genotype were examined. This included three biological replicates per genotype, with 20 images analyzed per replicate. The diameter of the endosomes was measured using the iTEM acquisition software (Olympus).

### 2.10 Scanning Electron Microscopy

iPSC-derived neurons were seeded at a density of 45,000 cells per well on glass coverslips (Thermo Fisher Scientific) placed in 24-well plates. After 20 days of maturation in CNM medium, wells were washed with PBS and fixed in 3% glutaraldehyde (Sigma) in 0.1 M cacodylate buffer (pH 7.3; Sigma) with 2% saccharose for 24 h at 4°C. Samples were washed with 0.1 M cacodylate buffer containing 2% saccharose and dehydrated in a series of increasing ethanol concentrations (30%, 50%, 70%, 96%, and 100% ethanol). They were then dried in a Critical Point Dryer (Bal-Tec CPD 030, LEICA). After that, samples were fixed on stubs and sputtered with gold using Q 150RS plus coater (Quorum Technologies). The samples were examined using TESCAN VEGA TS 5136 XM scanning electron microscope (TESCAN) at high vacuum mode, secondary electron detector mode, accelerating voltage of 15 kV at 3000X magnification.

For data analysis, approximately five projections per image were analyzed (>10 images per biological replicate). For each projection, the width and length of swellings, as well as the width and length of the shaft, were measured using ImageJ software. Minor shaft enlargements (<200% of shaft width), branching points, and enlargements at crossings were excluded. The number of swellings was then calculated per 100 µm of shaft length.

### 2.11 Live Imaging and axonal transport analyses

iPSC-derived neurons were seeded at a density of 60,000 cells per well in 8-well ibidi multichannel slide (Ibidi). Neurons were transduced at Day 20 with APP-GFP lentiviral particles (Flash Therapeutics) and with Bacmam 2.0 RFP-Rab5a (Thermo Fisher Scientific), both previously used in Feole *et al.* [20]. Particles were resuspended in CNM according to the validated multiplicity of infection (MOI; APP-GFP = MOI 5; RFP-Rab5a = MOI 10) and transduction units per milliliter (TU/mL) provided by the manufacturer. They were then incubated with cells for 24 hours, after which fresh CNM media was added. Transduction efficiency was monitored every 48 hours until the start of live-cell imaging experiments after 10 days of transduction (Day 30 of differentiation). Prior to acquisition, the CNM medium was replaced with artificial Cerebrospinal Fluid solution (aCSF) at pH 7.4 to avoid any interference from the phenol red contained in the original culture medium. Time-lapse recordings of axonal transport were performed in all genotypes for a total duration of 60 s, with varying frame rates depending on the cargo. Movement of APP-GFP vesicles was recorded at 2 frames per second (fps), while that of Rab5a vesicles was recorded at 1 fps. Recordings were obtained using a Zeiss LSM 780 microscope equipped with an LSM7 Live module, an incubation system to maintain cultures at 37 °C and 5% CO2, and acquisition was carried out with 63×/1.4 NA Plan Apochromat oil-immersion objective (Zeiss) to achieve maximum resolution.

For the following axonal transport analyses, the recorded movies were first processed in ImageJ and subsequently analyzed in Imaris v.10 (Oxford Instruments). Particle trajectories were identified using either the semi-automated “Spots” (APP-GFP) or “Surfaces” (RFP-Rab5a) tracking tool integrated with an Autoregressive Motion algorithm, for segmentation and thresholding of the APP and Rab5a particles. Input parameters included empirically determined XY diameters, maximum displacement distance, and maximum allowed gap size, tailored to the spatial and temporal resolution of each dataset. Key parameters used to assess axonal transport dynamics included track displacement (t*Dx*), defined as the spatial difference between the starting and ending *x*-coordinates of a particle (t*Dx*(t_1_, t_ₙ_) = P*x*(t_ₙ_) – P*x*(t_₁_) – where P indicates the position), and track duration (t*d*), indicating the total time a particle remained in motion. Tracks shorter than 10 seconds were excluded from further analysis. Based on average velocity, particles were classified as stationary or moving, and moving tracks were further categorized as anterograde or retrograde according to directionality. For APP-GFP, particles covering total distances > 250 nm were considered moving particles. If moving at average velocities of either ≥ 0.1 µm/s or -≤0.1 µm/s, they were considered fast anterograde and retrograde, respectively. All Rab5a-RFP particles, regardless of the speed, with a total track displacement ≥ 250 nm were considered in movement; otherwise, they were considered stationary. Track length was determined from the vectorial displacement (*x* and *y* dimensions) computed by the analysis software. Additionally, since movement of Rab5+ endosomes was analyzed using the “Surfaces” tool, the algorithm also allowed for the quantification of endosomal size over time, using the same time-lapse recordings acquired for axonal transport analysis.

### 2.12 Electrophysiology

iPSC-derived neurons were seeded at a density of 20,000 cells/well in the Poly-L-Ornithine/Laminin-coated 96-well MEA plates (Axion BioSystems). Each cell clone was seeded in 8 separate wells per individual run. Neurons were maintained in CNM for a period of 8 weeks. Spontaneous neuronal activity was recorded every other day for 5 minutes at 37 °C using the Axion Maestro system with the Middleman interface (Axion Biosystems). Raw electrophysiological data were processed using the Neural Metrics Tool in AxiS 2.4 software (Axion Biosystems) and exported for further analysis. Activity detection parameters were defined as follows: (a) an electrode was considered active if it recorded >1 spike per minute; (b) spike detection threshold was set at 6.5 standard deviations above the baseline noise; and (c) bursts were defined as a minimum of 5 spikes with an inter-spike interval no greater than 100 ms. To assess network maturation and activity, the following five parameters were analyzed: (1) weighted mean firing rate per well (average firing rate across active electrodes), (2) number of active electrodes per well, (3) percentage of spikes occurring within bursts, (4) number of bursts per well, and (5) synchrony index, representing the temporal correlation of firing across electrodes in a well.

### 2.13 Data analysis

Data are presented as mean ± standard error of the mean (SEM). Normality of data distribution was assessed prior to statistical testing. Parametric tests (t-test or one-way ANOVA) were applied to normally distributed data, while non-parametric tests (Mann–Whitney U test or Kruskal–Wallis test) were used for non-normally distributed data when comparing two or three groups, respectively. Statistical analyses were performed using GraphPad Prism version 9.0.0 for Windows (GraphPad Software; www.graphpad.com). A p-value ≤ 0.05 was considered statistically significant.

## 3. Results

### 3.1 The pathogenic SORLA p.Y1816C variant impairs SORLA protein maturation in iPSCs

In this study, we used isogenic human iPSC lines carrying either wild-type (WT) or heterozygous *SORL1* p.Y1816C variant, reflecting the genotype found in AD patients, hereafter referred to as knock-in (KI). We also included *SORL1* knock-out (KO) line. Both KI and KO iPSCs were generated via CRISPR/Cas9 genome editing [17] (**Figure 1A**). The pluripotent status of all lines was confirmed by characteristic colony morphology and stable expression of the pluripotency marker OCT4, the level of which remained unchanged following gene editing (**Figure 1B–D**). Furthermore, parental cell line exhibited a normal karyotype and was confirmed to carry the APOE3/3 genotype (**Figure S1A,B**). The presence of the correct point mutation in the *SORL1* gene was verified via Sanger sequencing (**Figure S1C**).

**Figure 1:**
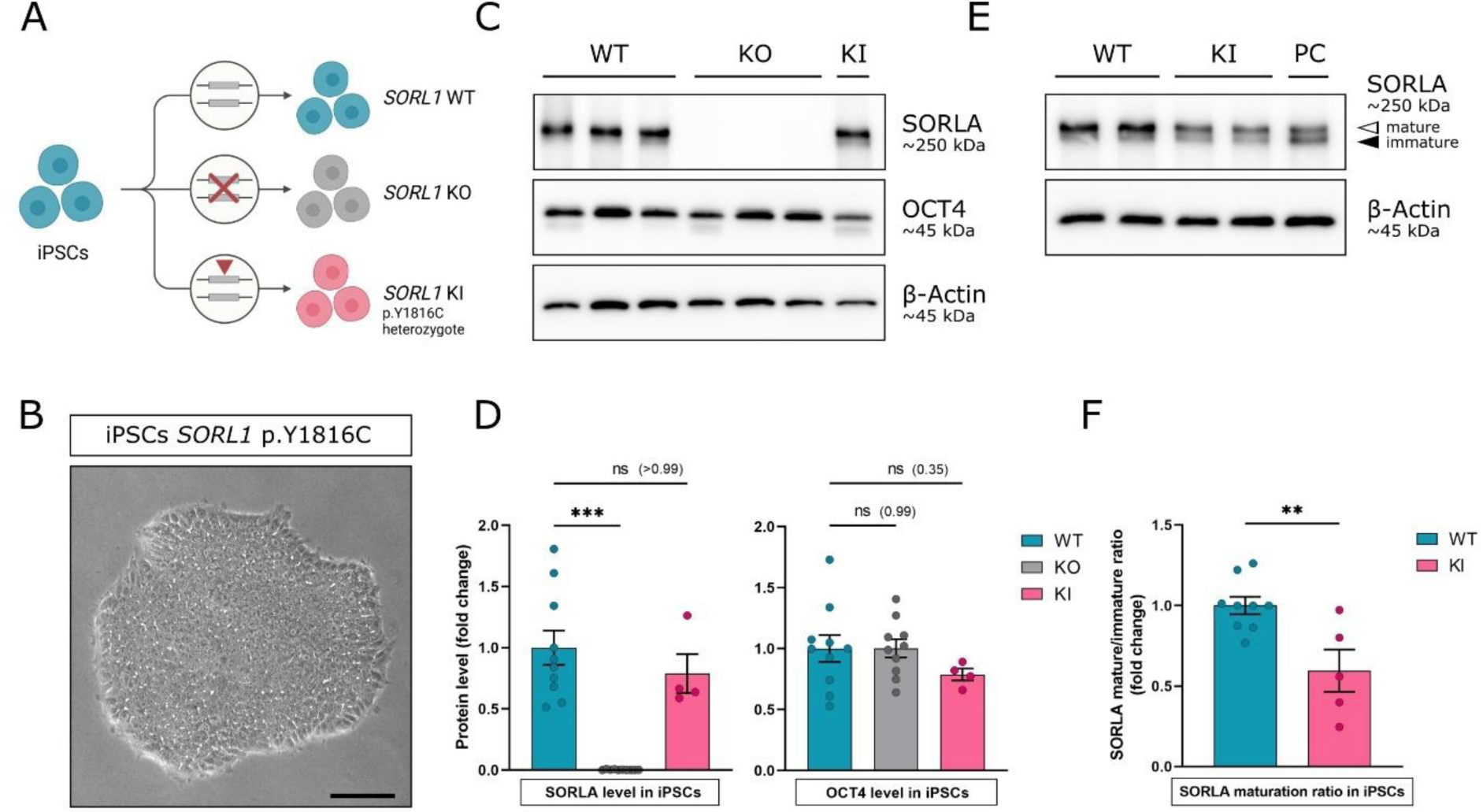
Characterization of isogenic iPSC lines carrying the *SORL1* p.Y1816C variant and analysis of SORLA protein maturation. **(A)** Schematic overview of the CRISPR/Cas9 gene editing strategy used to generate isogenic *SORL1* knock-in (KI; heterozygous p.Y1816C) and knock-out (KO) iPSC lines; **(B)** Representative colony morphology of pluripotent iPSCs carrying the *SORL1* p.Y1816C variant; **(C, D)** Western blot analysis on 10% PAGE gel showing expression and densitometric quantification of OCT4 and total SORLA across wild-type (WT), KI, and KO lines. All clones used in this study are shown; **(E)** Western blot on 6% PAGE gel visualizing the mature and immature forms of SORLA in WT and KI lines. PC – positive control cell line with compromised SORLA maturation [13]; **(F)** Densitometric quantification of the ratio of mature to immature SORLA forms; β-Actin was used as a loading control. Each dot represents one biological replicate (n ≥ 4). Data are presented as mean ± SEM. Statistical significance was assessed using one-way ANOVA followed by Dunnett’s Tukey’s multiple comparison test, Kruskal–Wallis followed by Dunn’s multiple comparison test or t-test; **p < 0.01, ***p < 0.001; Brightfield scale bar: 200 µm. A list of all experimental samples, including iPSC clones used for individual analyses and the number of biological replicates, is provided in **Table S2**

To evaluate the potential functional consequences of the p.Y1816C mutation, we next examined SORLA protein expression and maturation at the undifferentiated stem cell state of the cell lines. Given that SORLA undergoes essential post-translational processing required for its intracellular trafficking and activity, we analyzed both the immature and mature forms of the protein. While the total SORLA protein levels from expression of both alleles remained unchanged in KI cells compared to WT controls (**Figure 1C,D**), we observed a pronounced reduction in the mature form of SORLA in the KI line (**Figure 1E**). Further densitometric analysis and quantification of the ratio between mature and immature SORLA forms revealed a significant decrease, resulting in approximately a 40% reduction in SORLA maturation (**Figure 1F**). Overall, these initial data demonstrate that the pathogenic *SORL1* p.Y1816C variant does not affect the pluripotency of generated iPSCs but leads to impaired processing of the SORLA protein in iPSCs. Given that reduced levels of mature, functional SORLA have been linked to disrupted endosomal trafficking and increased AD risk [33], our initial findings prompted us to further explore pathogenic mechanisms by which the *SORL1* p.Y1816C variant may contribute to disease development at a cellular level, particularly in neuronal models.

### 3.2 The *SORL1* p.Y1816C variant reduces SORLA protein shedding in neurons and cerebral organoids

To further comprehensively assess the cellular consequences of the *SORL1* p.Y1816C mutation and SORLA deficiency in human neurons, we conducted parallel experiments using both 2D *NGN2*-induced neurons and 3D cerebral organoids derived from isogenic iPSC lines. This combined approach allowed us to validate findings across simplified 2D neurons and neuronal networks, as well as more complex 3D neural systems that mimic human brain architecture and cellular interactions [30,31].

For the generation of 2D neurons, we used a doxycycline-inducible *NGN2* overexpression system, as previously described [28,29]. Neuronal differentiation was monitored at three time points, days 20, 30, and 40, to track developmental progression (**Figure 2A**). All iPSC lines successfully differentiated into neurons, forming characteristic 2D neuronal networks and expressing canonical neuronal markers including TUJ1, NeuN, MAP2, and synaptophysin (SYP) (**Figure 2B**). Subsequent extensive molecular characterization confirmed robust neuronal identity across all genotypes. While some variability in differentiation timing of gene or protein expression was observed between biological replicates, the expression of lineage- and function-specific markers followed consistent patterns across experiments, irrespective of SORL1 WT, KO, or KI genotypes. Specifically, we analyzed neural stem cell markers (SOX2, SOX1, PAX6), neuronal markers (DCX, TUJ1, MAP2, NeuN), synaptic markers (SYN1, SYP, PSD95), and synaptic receptor components (GLUA, GLUN1, GLUN2A, GLUN2B, TRKB, TRKC) at both RNA and protein levels, providing strong evidence of neuronal specification (**Figure 2C and S2A,B**).

**Figure 2:**
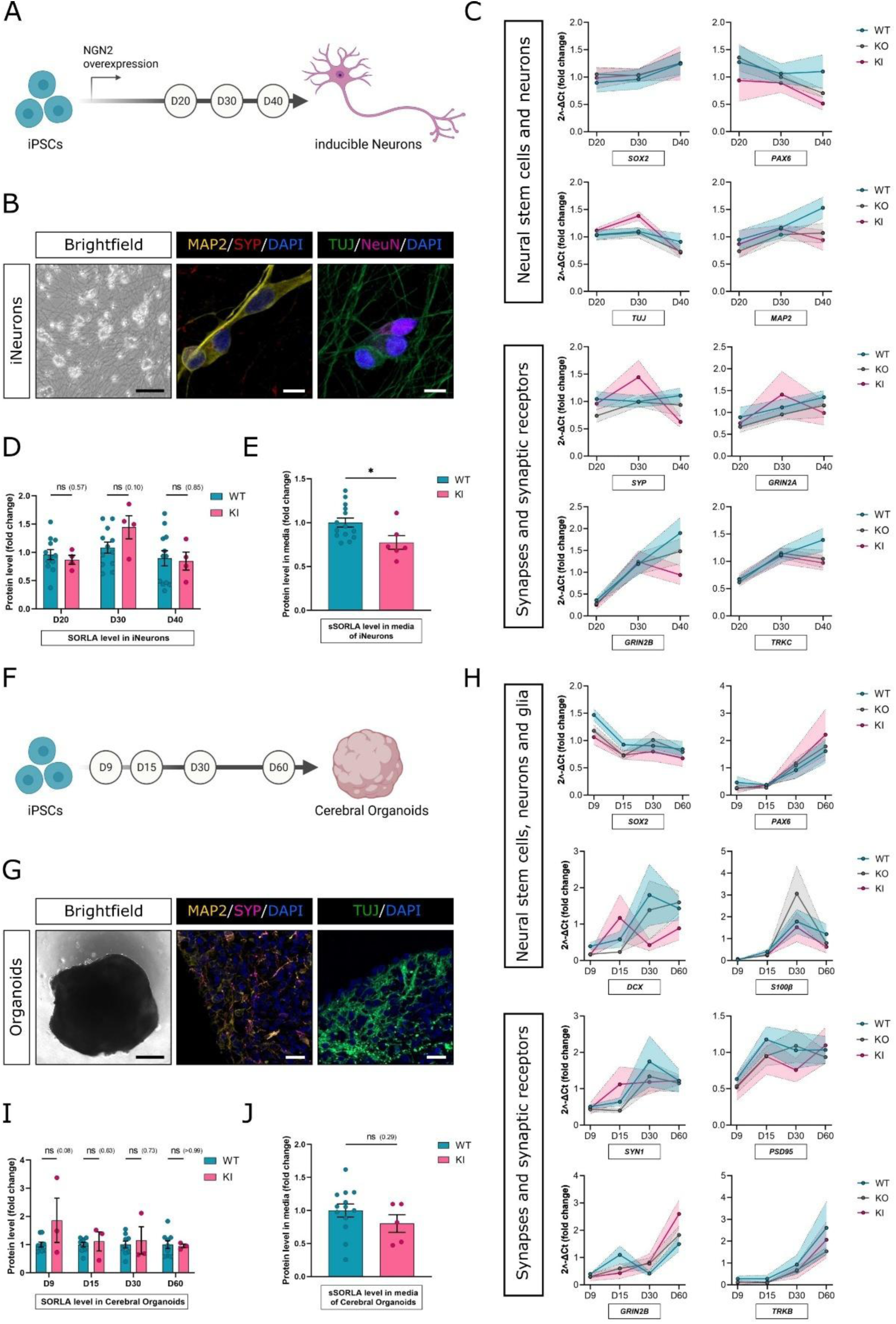
*SORL1* p.Y1816C variant affects SORLA shedding in 2D and 3D human models. **(A)** Schematic of the differentiation protocol for generating 2D neurons using a doxycycline-inducible NGN2 overexpression system and sample collection; **(B)** Representative images showing neuronal differentiation of iPSC lines, with expression of TUJ1, NeuN, MAP2, and synaptophysin (SYP); **(C)** Gene expression analysis of neural stem cell (SOX2, PAX6), neuronal (TUJ1, MAP2), and synaptic markers and receptors (SYP, GLUN2A, GLUN2B, TRKC); **(D)** Western blot analysis showing levels of intracellular SORLA during neuronal differentiation in WT and KI lines; **(E)** ELISA analysis of sSORLA levels in the culture medium of WT- and KI-derived neurons at D30; **(F)** Schematic of the protocol used for generating 3D cerebral organoids and timeline of sample collection; **(G)** Representative images showing day 60 cerebral organoids with expression of mature neuronal markers (MAP2, SYP, NeuN); **(H)** Gene expression profiles of progenitor (*SOX2, PAX6*), neuronal (*DCX*), synaptic (*SYN1, PSD96, GRIN2B, TRKB*), and glial (*S100β*) markers across developmental progression of WT, KO, and KI lines. **(I)** Total SORLA protein expression compared between genotypes during organoid differentiation. **(J)** ELISA analysis of sSORLA in day 60 organoid culture medium. Each dot in (C) and (H) represents the average 2˄-ΔCt; each dot in (D), (E), (I), and (J) represents one biological replicate (n ≥ 3). Data are presented as mean ± SEM. Statistical significance was assessed using Mann-Whitney test or t-test; *p < 0.05 was determined using an unpaired statistical test. Brightfield scale bar: 50 µm (B), 250 µm (G); Immunofluorescence scale bar: 10 µm (B), 20 µm (G). A list of all experimental samples, including iPSC clones used for individual analyses and the number of biological replicates, is provided in **Table S2**

Importantly, we also assessed the expression and processing of SORLA during neuronal differentiation. Results show that intracellular SORLA protein levels remained comparable between wild-type and p.Y1816C mutant neurons (**Figure 2D**). However, quantification of shedded sSORLA in the culture medium at day 30 by ELISA revealed a significant reduction in mutant neurons compared to controls (**Figure 2E, p < 0.05)**. This decrease mirrors previously reported reductions in sSORLA levels in cerebrospinal fluid from human AD patients carrying *SORL1* mutations, further supporting the relevance of our *in vitro* model [29].

In parallel, 3D cerebral organoids were generated using a modified Lancaster protocol [30,31] and collected at four developmental stages (days 9, 15, 30, and 60; **Figure 2F**). By day 60, the organoids displayed typical morphology and regional organization and expressed mature neuronal markers, including MAP2, SYP, and NeuN (**Figure 2G**). Further analyses confirmed that the expression of pluripotency markers progressively declined during differentiation in 3D organoids, while progenitor, neuronal, synaptic, and glial markers followed expected developmental trajectories. Importantly, no consistent differences were observed between control, SORLA KO, or *SORL1* p.Y1816C variant lines when assessing both qPCR and Western blot results. (**Figure 2H and S2C,D**). Notably, similarly to 2D neurons, total levels of SORLA protein detected in cerebral organoids over the course of differentiation remained comparable between analyzed cell lines (**Figure 2I**). However, sSORLA levels measured by ELISA in the day 60 culture medium showed a decreasing trend in mutant lines, consistent with 2D neuron data, though the difference did not reach statistical significance (**Figure 2J**). Two additional biological replicates analyzed by Western blot revealed a similar trend, with reduced sSORLA levels in KI-derived organoids compared to WT controls (**Figure S2E,F**). Together, these findings demonstrate that the *SORL1* p.Y1816C variant impairs SORLA maturation in iPSC cultures and shedding in both 2D neuronal cultures and more physiologically relevant 3D cerebral organoids, highlighting a consistent defect in SORLA processing across complementary human-based models expressing the pathogenic mutation endogenously.

### 3.3 p.Y1816C induces enlargement of early endosomes in both 2D neurons and 3D organoids

Thus far, our data have demonstrated that the *SORL1* p.Y1816C variant alters the biology of the SORLA protein itself, leading to impaired maturation and reduced shedding. To further assess the impact of this variant on neuronal physiology, we next investigated the effect of reduced SORLA activity on endosomal biology, a process previously associated with SORLA dysfunction. Indeed, we and others have reported that enlarged early endosomes represent a pathological hallmark of impaired SORLA function [16,29,34].

To evaluate this, we selected day 30 (D30) for 2D neuronal cultures and day 60 (D60) for 3D cerebral organoids as representative time points for our confocal microscopy analysis. We first performed Immunofluorescence staining for EEA1, a marker of early endosomes, in both models. Representative images from confocal microscopy (**Figure 3A**) confirmed the previously observed phenotype and revealed visibly enlarged EEA1-positive endosomes in both SORLA KI and KO neurons compared to WT controls. A similar pattern was observed in 3D cerebral organoids, where KI and KO lines exhibited a greater proportion of visibly enlarged endosomes than WT. Quantitative image analysis of 2D neurons further confirmed these observations. Specifically, while endosomes of various sizes were present across all genotypes (**Figure S3A**), the proportion of standard-sized endosomes (≤0.35 µm^3^) in WT neurons was approximately 92.2%, with only 7.8% falling into the category of enlarged endosomes (>0.35 µm²). In contrast, neurons derived from KI and KO iPSCs showed a substantial increase in larger endosomes, with 12.9% classified as enlarged and only 88.1% as standard-sized (**Figure 3B**). The distribution of the size of enlarged endosomes across genotypes is then shown in **Figure 3C**. Notably, when comparing the total number of enlarged endosomes across four biological replicates and a total of 84 individual images of neuronal cultures across all three genotypes, we detected a >1.5-fold increase in KI neurons (n = 195 endosomes) and a >3.6-fold increase in KO neurons (n = 456 endosomes) compared to WT (n = 124 endosomes) (**Figure 3D**). All these data confirm that neurons carrying the AD-causing mutation in *SORL1* have indeed a higher proportion of enlarged endosomes, as reported before [16,29,34,35].

**Figure 3.**
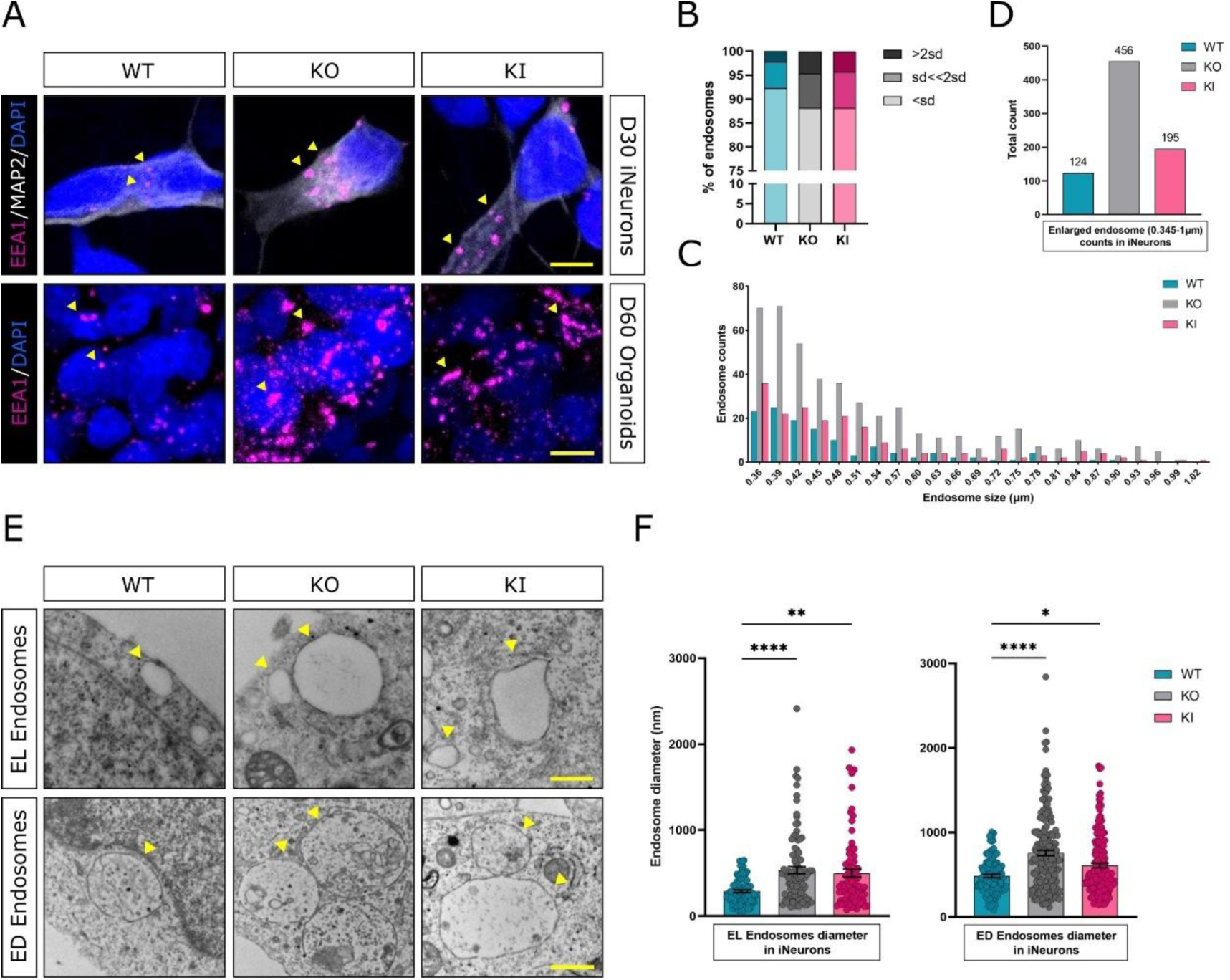
Enlargement and altered distribution of early endosomes in *SORL1* p.Y1816C and KO neurons and organoids. **(A)** Representative immunofluorescence images showing EEA1-positive early endosomes in WT, SORLA KO, and KI 2D neurons at day 30 and 3D cerebral organoids at day 60; **(B)** Quantification of endosome size categories in 2D neurons, showing the proportion of standard-sized (≤0.35 µm²) and enlarged (>0.35 µm²) EEA1-positive endosomes across genotypes. Visualized are endosomes percentages of normal, enlarged, and large categories as classified by the standard deviation, normal < sd, enlarged sd>2sd, and large >2sd; **(C)** Distribution of EEA1-positive endosomes with size >0.35 µm² in 2D neurons derived from WT, KI, and KO lines. Size distribution of all endosomes is shown in **Supplementary Figure 3A; (D)** Total number of enlarged endosomes (>0.35 µm²) detected per condition across four biological replicates and over 12 images per genotype in total. **(E)** Representative transmission electron micrographs (TEM) of electron-lucent (EL) and electron-dense (ED) endosomes in WT, KI, and KO neurons; **(F)** Quantification of the diameter of EL and ED endosomes measured from TEM images. Each dot represents one endosome (biological replicates: n ≥ 3). Data are presented as mean ± SEM. Statistical significance was assessed using Kruskal–Wallis followed by Dunn’s multiple comparison test; ****p < 0.001, **p < 0.01, *p < 0.05. Immunofluorescence scale bar: 5 µm (iNs) 10 µm (COs). TEM scale bar: 500 nm. A list of all experimental samples, including iPSC clones used for individual analyses and the number of biological replicates, is provided in **Table S2.**

Importantly, although the enlargement of endosomes in SORLA KO neurons has been previously described using immunocytochemistry, a detailed ultrastructural analysis by transmission electron microscopy (TEM) has been lacking. We therefore performed TEM on three biological replicates of differentiated neurons at D20 across genotypes to examine the size, content, and distribution of endosomal structures across the analyzed genotypes. Based on existing literature [36], we identified and classified ultrastructural features of the endolysosomal system, including endocytosis and clathrin-coated pits, electron-lucent (EL) endosomes, electron-dense (ED) endosomes, and vesicles undergoing exocytosis (**Figure S3B**). Notably, EL and ED endosomes were present in WT neurons in a range of sizes, with ED endosomes generally being larger, prompting us to analyze their dimensions separately. Results, as depicted on representative TEM micrographs (**Figure 3E**) and supported by quantitative analysis (**Figure 3F**), showed substantial differences in the size of both EL and ED endosomes across genotypes, with both EL and ED endosomes being significantly enlarged in KO neurons and KI neurons compared to WT.

Interestingly, beyond size, we also observed spatial clustering of enlarged endosomes in KI and KO neurons. In these cells, multiple enlarged endosomes frequently appeared clustered within the soma, often visible within a single TEM field (**Figure 3D and S3C**). In contrast, WT neurons displayed more evenly distributed endosomes. We did not observe any abnormal fusion events between endosomes in any of the genotypes. Taken together, these results show that the *SORL1* p.Y1816C variant causes endosomal enlargement in 2D neurons and 3D cerebral organoids, closely resembling the phenotype observed in SORLA KO models. Furthermore, characterization of the ultrastructure of enlarged endosomes in SORLA mutant human neurons revealed both morphological changes and clustering behavior that may reflect defects in endosomal trafficking and intracellular positioning, processes essential for normal neuronal function [37,38].

### 3.4 p.Y1816C leads to APP accumulation in early endosomes, increased secretion of Aβ40 and Aβ42, and enhanced amyloid β cluster deposition in cerebral organoids

Thus far, we have demonstrated that the *SORL1* p.Y1816C mutation, associated with familial AD, leads to the enlargement of early endosomes in both 2D neurons and 3D cerebral organoids. Importantly, we and others have previously shown that in SORLA-deficient cellular models or iPSC-derived neurons, such endosomal enlargement leads to APP accumulation within endosomes and increased cleavage of APP into Aβ40 and Aβ42 peptides [16,39,40]. However, it remained unclear whether these downstream effects also occur in human neurons carrying a point mutation in *SORL1*, and whether this phenotype is preserved in more complex 3D systems. Thus, to address these questions, we first analyzed 2D neuronal cultures and assessed co-localization between APP and EEA1 using the proximity ligation assay (PLA). This technique detects protein–protein proximity (≤40 nm), producing a fluorescent signal when two targets, here APP and EEA1, are spatially close, suggesting endosomal co-localization. Results, as shown in **Figure 4A**, confirmed that PLA signals were detectable in all tested conditions, but the number of APP–EEA1 foci was markedly elevated in KI and KO neurons compared to WT. Quantitative analysis revealed a ∼3-fold increase in PLA signal per nucleus in both KI and KO neurons relative to WT (**Figure 4B**), indicating enhanced APP retention within early endosomes. Additionally, as detected by ELISA, both KI and KO neurons exhibited a trend toward increased Aβ40 and Aβ42 secretion compared to WT (**Figure 4C**), consistent with prior observations in SORLA-deficient neurons [16,40].

**Figure 4.**
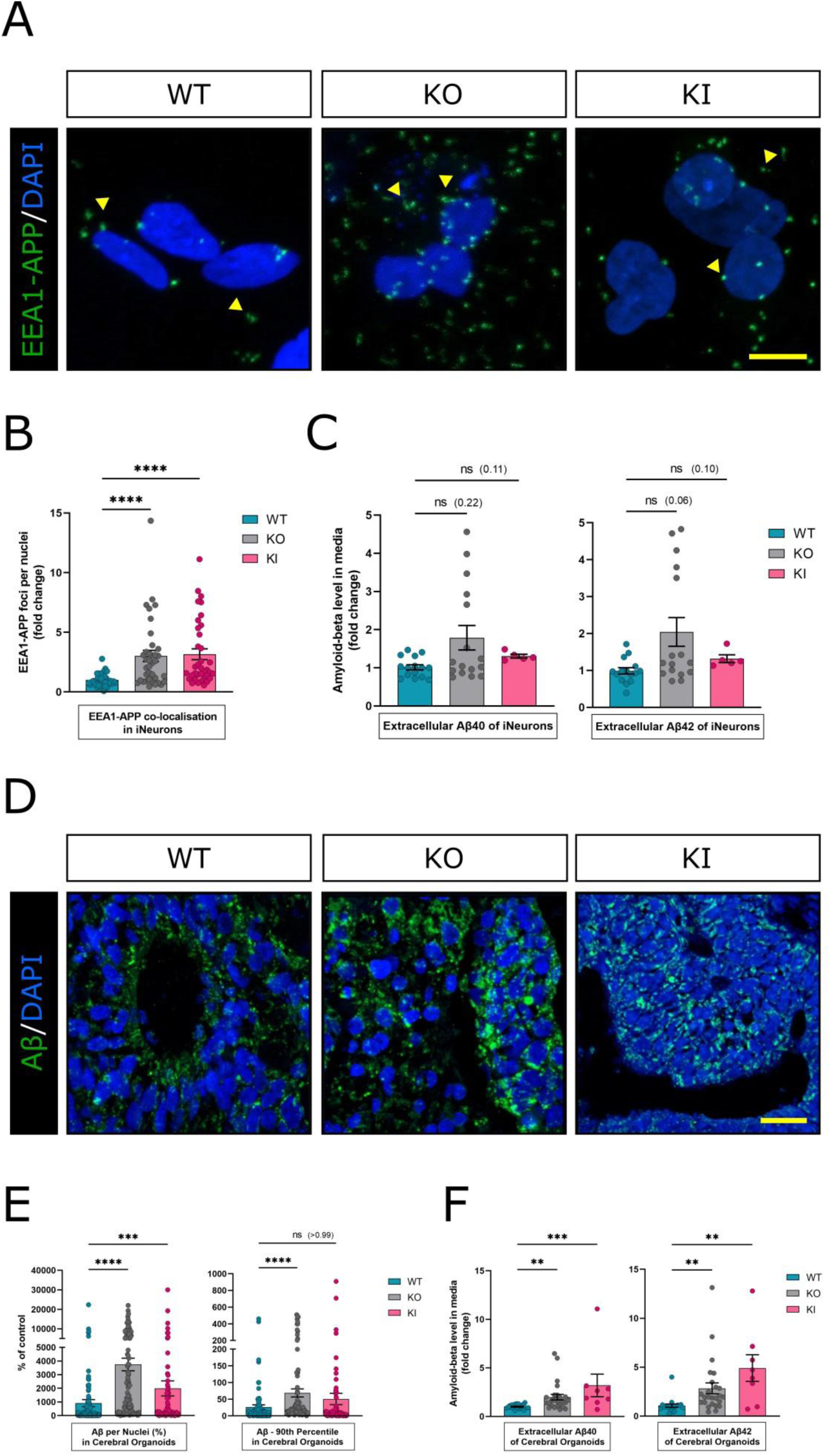
The *SORL1* p.Y1816C mutation promotes APP accumulation in early endosomes and increases Aβ secretion and deposition in neurons and cerebral organoids. **(A)** Representative images from proximity ligation assay (PLA) detecting APP–EEA1 co-localization in day 20 2D neurons derived from WT, SORLA KI, and KO iPSC lines. **(B)** Quantification of PLA signal per nucleus; **(C)** ELISA measurement of secreted Aβ40 and Aβ42 levels in 2D neuronal culture media; **(D)** Representative immunofluorescence images of sections from D60 cerebral organoids stained for amyloid-β (Aβ) **(E)** Quantification of Aβ signal in organoid sections, including: (i) total Aβ signal per cell (normalized to nuclear volume) and (ii) number of large Aβ clusters (defined as 90th percentile of particle size). The average signal intensity of the control samples was used as a reference for comparing all analyzed data; **(F)** ELISA measurements of secreted Aβ40 and Aβ42 to organoid cell culture media. Each dot represents one biological replicate (n ≥ 3). Data represent mean ± SEM. Statistical significance was assessed using Kruskal–Wallis followed by Dunn’s multiple comparison test; *p < 0.05, **p < 0.01, ***p < 0.001, ****p < 0.0001. Immunofluorescence scale bar: 10 µm (A) 25 µm (D). A list of all experimental samples, including iPSC clones used for individual analyses and the number of biological replicates, is provided in **Table S2**

Importantly, to determine whether this phenotype extends to 3D models, we next examined Aβ and Aβ40/42 peptides in our cerebral organoids. We previously demonstrated that familial AD mutations in presenilin genes lead to Aβ accumulation in organoids and altered secretion of Aβ peptides [31]. Applying a similar workflow, we first stained 200-µm-thick sections for Aβ and observed increased Aβ signal in both KI and KO organoids compared to WT (**Figure 4D**). Image analysis quantification further supported these findings. We specifically quantified i) the overall quantity of Aβ signal per cell (assessed by the volume of Aβ particles normalized to the volume of cell nuclei per image; **Figure 4E**), ii) changes in the overall size of all Aβ particles (visualized as the median volume of Aβ; **Figure S4A**), and iii) specifically the number of large Aβ clusters (quantified as the 90^th^ percentile of Aβ particle size; **Figure 4E**). These analyses revealed that, although the overall size distribution of Aβ particles was not significantly different across genotypes, both KI and KO organoids exhibited a significantly higher amount of Aβ per cell, and a trend toward an increased number of large Aβ clusters (**Figure 4E**). Finally, ELISA of organoid-conditioned media revealed a significant increase in secreted Aβ40 and Aβ42 levels in both KO and KI organoids compared to WT (**Figure 4F**). Importantly, APP mRNA expression levels remained comparable across WT, KI, and KO lines in both neuronal and organoid models (**Figure S4B**), suggesting that the observed phenotypes are not due to transcriptional changes but rather reflect post-translational dysregulation of APP trafficking and processing. Together, these results demonstrate that the *SORL1* p.Y1816C variant promotes APP retention in early endosomes, leading to increased Aβ production and accumulation, contributing to amyloid pathology in both 2D and 3D human neuronal models.

### 3.5 Altered SORLA increases neuritic swelling and deregulates endosomal transport along axons

Thus far, our analyses have focused primarily on the soma of neurons, where we observed endosomal enlargement, defined their ultrastructure via TEM, and demonstrated APP accumulation within these compartments. However, neurons are highly polarized cells, and their proper function depends on efficient intracellular transport not only within the soma but also along axons and dendrites, particularly for the delivery of materials to and from synaptic terminals [21]. Interestingly, during our TEM analyses of neurons with dysfunctional SORLA, we observed a cross-section of a neuronal axon exhibiting pronounced swelling, within which we identified an enlarged endosome (**Figure 5A**). Importantly, axonal swellings are strongly associated with disrupted axonal transport and serve as early indicators of neuronal dysfunction and neurodegeneration in various disease concepts, including AD [2,41]. Moreover, prior studies on iPSC-derived AD neurons carrying APP mutations have implicated impaired axonal transport as a central mechanism contributing to disease pathology [2,20,42]. Thus, this observation prompted us to systematically investigate the effects of SORLA deficiency and the *SORL1* p.Y1816C mutation on axonal morphology and transport dynamics.

**Figure 5:**
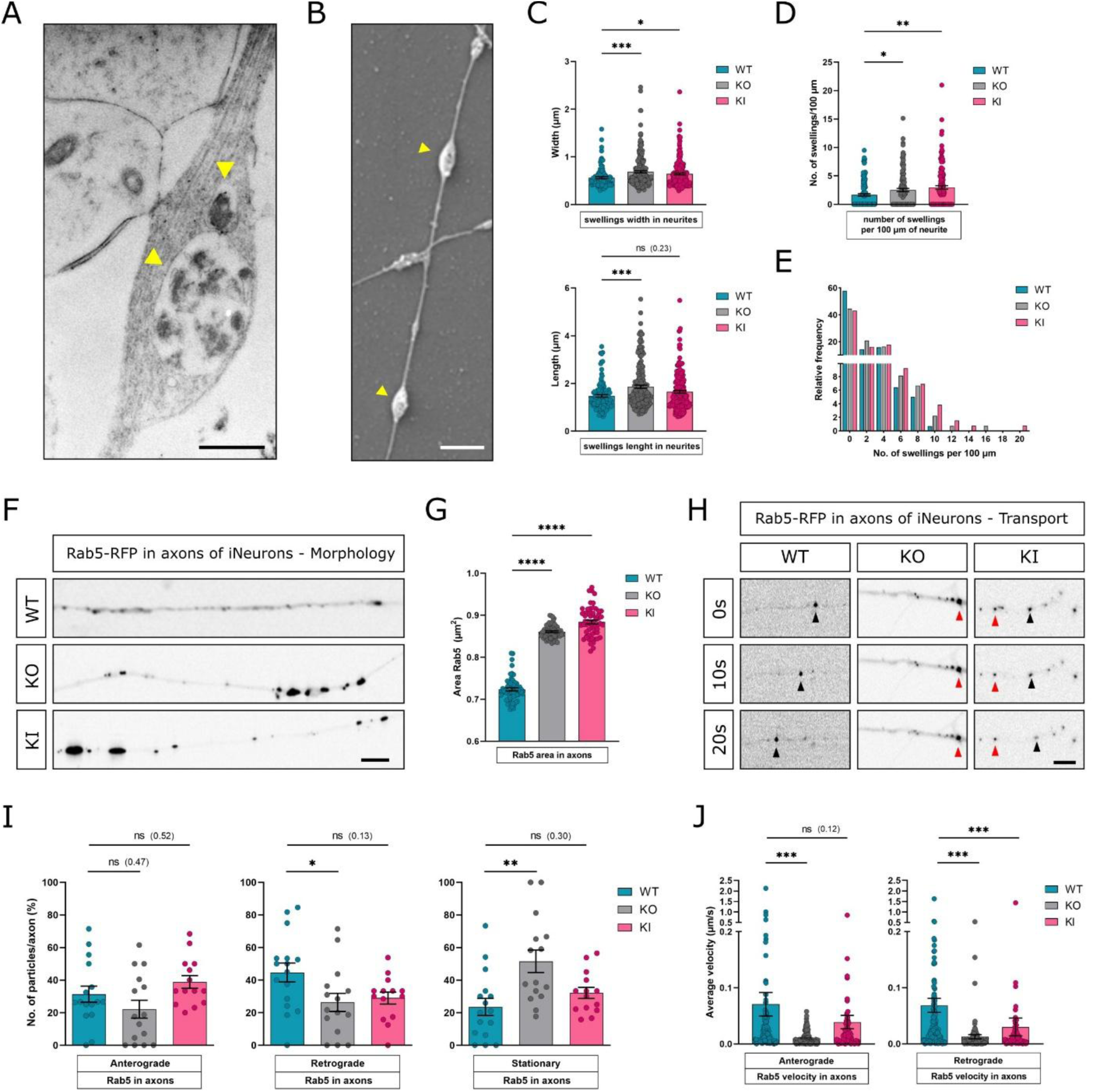
Altered SORLA increases neuritic swelling and deregulates endosomal transport along axons. **(A)** Transmission electron microscopy (TEM) micrograph showing a cross-section of a neuritic swelling containing an enlarged endosome; **(B)** Representative scanning electron microscopy (SEM) image of differentiated KI neurons displaying neuritic swellings; **(C)** Quantification of swelling width and length across genotypes; **(D)** Number of swellings per 100 µm of neurite in WT, KI, and KO neurons; **(E)** Distribution of the number of swellings per 100 µm segment of neurite; **(F)** Live-cell images of Rab5a-RFP signal in individual axons of WT, KI, and KO neurons at D30; **(G)** Quantification of Rab5+ signal area along axons; **(H)** Representative pictures showing Rab5+ endosome movement along axons for each genotype. Black arrows point to moving particles, red arrows point to stationary particles; **(I)** Quantification of axonal transport directionality and proportion of stationary Rab5+ particles; **(J)** Quantification of the velocity of anterograde and retrograde Rab5+ particle movement over a 60-second imaging period. In C and D, each dot represents a swelling. A total of 90 projections were analyzed from three biological replicates per genotype; In G, I, and J, each dot represents the mean of Rab5+ endosomes per time point (G), one axon (I), a single endosome (J). All the data were measured in three independent biological replicates. Data represent mean ± SEM. Statistical significance was assessed using one-way ANOVA followed by Dunnett’s Tukey’s multiple comparison test or Kruskal–Wallis followed by Dunn’s multiple comparison test or t-test; *p < 0.05, **p < 0.01, ***p < 0.001, ****p < 0.0001. Scale bar on a representative image of axon: 20 µm. TEM scale bar: 500 nm. SEM scale bar: 2 µm. A list of all experimental samples, including iPSC clones used for individual analyses and the number of biological replicates, is provided in **Table S2**

We began our analyses by quantifying neuritic swellings in 2D neurons derived from WT, KO, and KI lines. To assess these changes in our model, we performed scanning electron microscopy (SEM) on differentiated WT, KI, and KO neurons, using previously established methodology in Pozo *et al.* [41]. SEM imaging clearly revealed neuritic swellings across all genotypes, allowing for the quantification of multiple morphological parameters (**Figure 5B**). Our analysis showed that SORLA deficiency and mutation both affect neuritic swelling parameters. Specifically, the width of neuritic swellings was significantly increased in both KO and KI neurons compared to WT (**Figure 5C**). The length of individual swellings was also significantly longer in KO neurons and showed a trend toward an increase in KI neurons. When we calculated the total number of swellings per 100 µm of neurite, we observed a significantly increased swellings in both KO and KI neurons compared to WT (**Figure 5D**). Importantly, when we visualized the distribution of swellings per 100 µm, WT neurites generally contained fewer swellings, while KI and KO neurites showed a shift toward a higher frequency of swellings (**Figure 5E**). Notably, neurites containing more than 10 swellings per 100 µm were rare and exclusively observed in KO and KI neurons, but never in WT. These findings suggest that SORLA dysfunction, either through loss or mutation, leads to structural abnormalities along axons, potentially indicating compromised vesicular transport and endosomal trafficking, which are critical for maintaining neuronal health and polarity.

Thus, we next sought to analyze endosomal trafficking along axons in more detail. To achieve this, we utilized our system of inducible neurons, which were differentiated until day 20 (D20). Subsequently, we labeled neuronal cultures with CellLight Early Endosomes-RFP, a BacMam 2.0 reagent that expresses red fluorescent protein (RFP)-tagged Rab5a, allowing for the live-cell visualization of early endosomes (hereafter referred to as Rab5+). On D30, we performed live-cell imaging of individual axons in WT, KI, and KO neurons and quantified their axonal transport dynamics. As shown in **Figure 5F** and quantified in **Figure 5G and S5A**, Rab5+ signals already displayed distinct patterns across genotypes. In WT neurons, Rab5+ endosomes appeared as small puncta distributed evenly along the axon. In a strong contrast, KI and especially KO neurons exhibited large Rab5+ clusters, indicating disrupted endosomal distribution. Importantly, changes in the area of Rab5⁺ endosomes were observed only when comparing WT, KO, and KI genotypes, but not across time within the same genotype, suggesting that once endosomes are enlarged, their size remains stable over at least a 60-second period (**Figure S5A**).

Interestingly, quantitative analysis of Rab5+ endosomal transport revealed clear genotype-dependent differences in directionality and motility (**Figure 5H, I, and S5B**). In WT neurons, Rab5+ particles displayed typical transport behavior, with ∼45% moving retrogradely, 32% anterogradely, and 23% remaining stationary, consistent with previous reports [20]. In KI neurons carrying the p.Y1816C variant, this distribution was altered, showing a reduced proportion of retrograde transport and a similar proportion of stationary particles, with modest differences in the anterograde fraction. The effect was more pronounced in SORLA KO neurons, where over 50% of Rab5+ particles remained stationary and fewer particles moved in either direction.

We next focused on the dynamic properties of the moving particles over a 60-second imaging period, excluding particles lower than 0.001 µm/s. Importantly, we found that velocities of both anterograde and retrograde Rab5+ particles were significantly reduced in KO neurons compared to WT, and trended lower in KI neurons (**Figure 5J**). The total distance traveled by individual endosomes, referred to as track length, was also significantly shorter in both KI and KO neurons (**Figure S5C**). Taken together, these results indicate that the *SORL1* p.Y1816C mutation, and more severely, complete SORLA loss, compromise both the movement and spatial distribution of Rab5+ endosomes in axons, highlighting a key role for SORLA in axonal endosomal trafficking.

### 3.6 SORLA alterations affect the axonal transport of APP

Given the striking disruption of endosomal distribution and trafficking along axons in our previous experiments, we next investigated whether axonal transport of APP, a critical protein in AD pathology, is similarly affected. Earlier studies, including ours, have demonstrated that familial AD mutations, such as the Swedish APP variant, impair retrograde motor function and broadly disturb axonal transport dynamics [20]. However, whether the SORLA protein has any effect on the axonal transport of APP remained unaddressed. Thus, to assess APP trafficking in this context, we used lentiviral constructs expressing GFP-tagged wild-type APP (APP-GFP; previously used in [20] as APPwt-GFP) and performed live imaging of axonal transport in 2D inducible neurons derived from WT, KO, and KI SORLA lines. As shown in **Figure 6A**, in contrast to the clustered Rab5+ signal observed previously, the APP-GFP signal appeared more uniformly distributed along axons.

**Figure 6.**
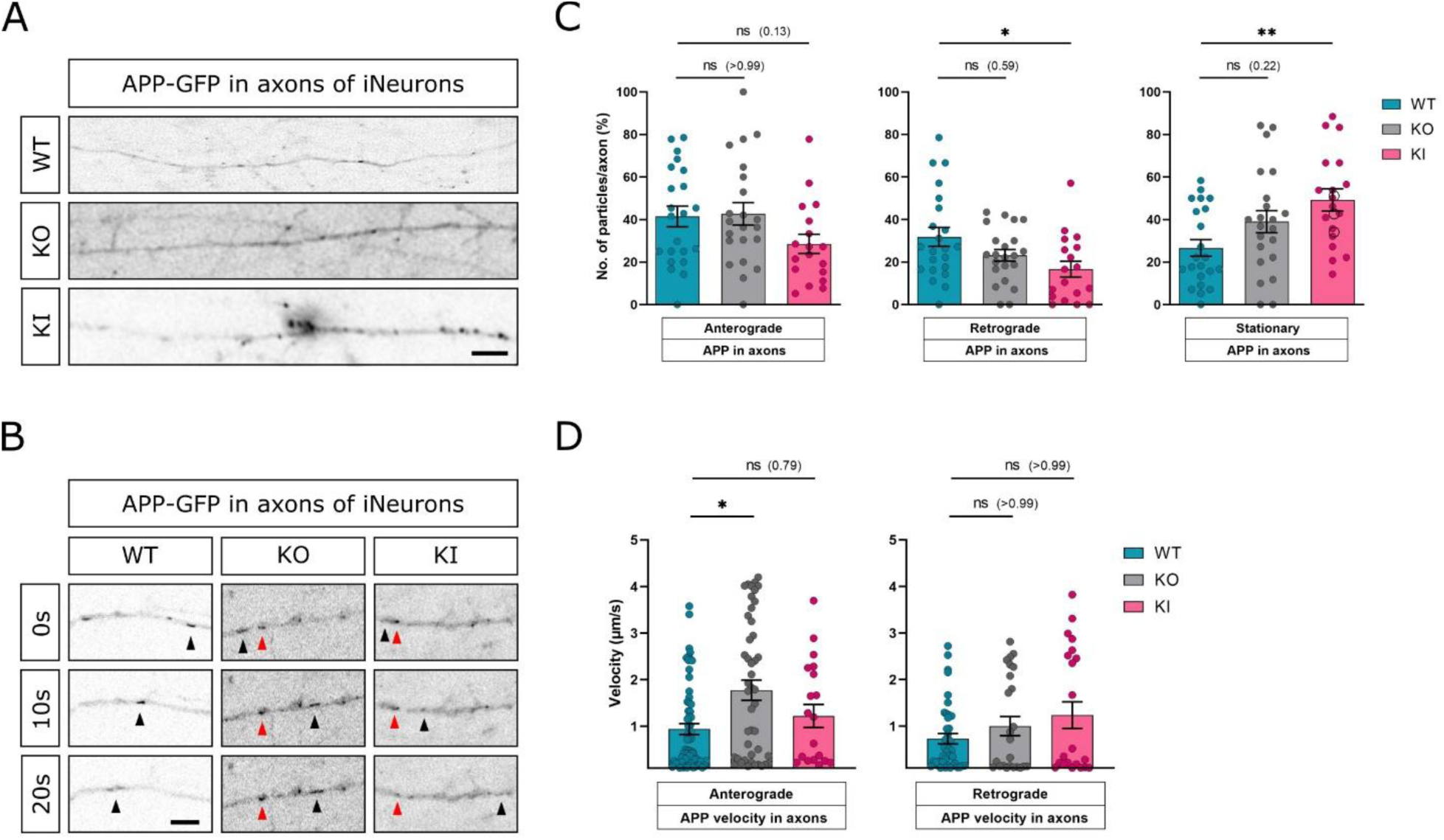
SORLA alterations impair axonal transport of APP in human neurons. **(A)** Representative still frames from live imaging of 2D neurons expressing APP-GFP, showing axonal localization of GFP signal in WT, KI, and KO neurons; **(B)** Representative pictures illustrating APP-GFP particle movement along axons in WT, KI, and KO neurons. Black arrows point to moving particles, Red arrows point to stationary particles; **(C)** Quantification of APP-GFP transport parameters showing the proportion of particles moving anterogradely, retrogradely, or remaining stationary; **(D)** Velocity of fast-moving APP-GFP particles (≥0.1 μm/s) in anterograde and retrograde directions across genotypes. Each dot represents one axon (C) or a moving APP particle (D and E). Data were measured in three independent biological replicates. Data represent mean ± SEM. Statistical significance was assessed using one-way ANOVA followed by Dunnett’s Tukey’s multiple comparison test, Kruskal–Wallis followed by Dunn’s multiple comparison test or t-test; *p < 0.05, **p < 0.01. Brightfield scale bar: 20 µm. Scale bar on a representative image of axon: 20 µm. A list of all experimental samples, including iPSC clones used for individual analyses and the number of biological replicates, is provided in **Table S2.**

However, tracking analyses revealed significant alterations in APP transport dynamics. Both *SORL1* KO and KI neurons showed a marked increase in the proportion of stationary APP-GFP particles compared to WT neurons. Although anterograde and retrograde transport remained detectable and followed a similar directional pattern across genotypes, the overall number of moving APP particles was reduced in KO and KI neurons, indicating a shift toward particle stalling. Representative images are shown in **Figure 6B**, with quantification in **Figure 6C** and **Figure S6A**. Moreover, the track length, which represents the total distance traveled by individual APP-GFP particles, was significantly shorter in KO and KI neurons compared to WT, suggesting impaired transport range and reduced mobility within axons (**Figure S6B**). Finally, analysis of the fast-moving subset of APP-GFP particles (≥0.1 μm/s) revealed that both anterograde and retrograde velocities were significantly elevated or showed an upward trend in KO and KI neurons compared to WT (**Figure 6D**). These findings suggest that SORLA dysfunction not only increases particle stalling but also alters the kinetics of moving APP vesicles, potentially reflecting compensatory changes in motor-cargo dynamics. Together with the observed impairments in Rab5⁺ endosomal transport, our results highlight a broader and previously underappreciated role for SORLA in regulating axonal transport and maintaining neuronal homeostasis in the context of AD pathology.

### 3.7 Electrophysiological measurements show hyperexcitable neurons with SORLA KO and *SORL1* p.Y1816C variant

Finally, to investigate the functional consequences of SORLA deficiency and the *SORL1* p.Y1816C mutation on neuronal activity, we performed long-term electrophysiological recordings using microelectrode array (MEA) technology on 2D differentiated neurons from D14 to D56. We assessed both individual electrode and global network parameters across WT, KO, and KI neuronal cultures. For individual electrode activity (**Figure 7A**), we analyzed the weighted mean firing rate, which showed a significant increase in KI neurons and a strong upward trend in KO neurons compared to WT. Similarly, the number of active electrodes was significantly elevated in both KI and KO neurons, suggesting increased spontaneous activity. Next, we evaluated network-level properties (**Figure 7B**). The burst percentage, a measure of neuronal firing maturation, was significantly higher in both KI and KO neurons, and we also found a significant increase in the number of bursting electrodes, indicating a maturation of electrophysiological properties of the neuronal cultures. Finally, the synchrony index, reflecting the degree of temporal coordination between electrodes and network engagement, was also significantly elevated in both mutant conditions.

**Figure 7:**
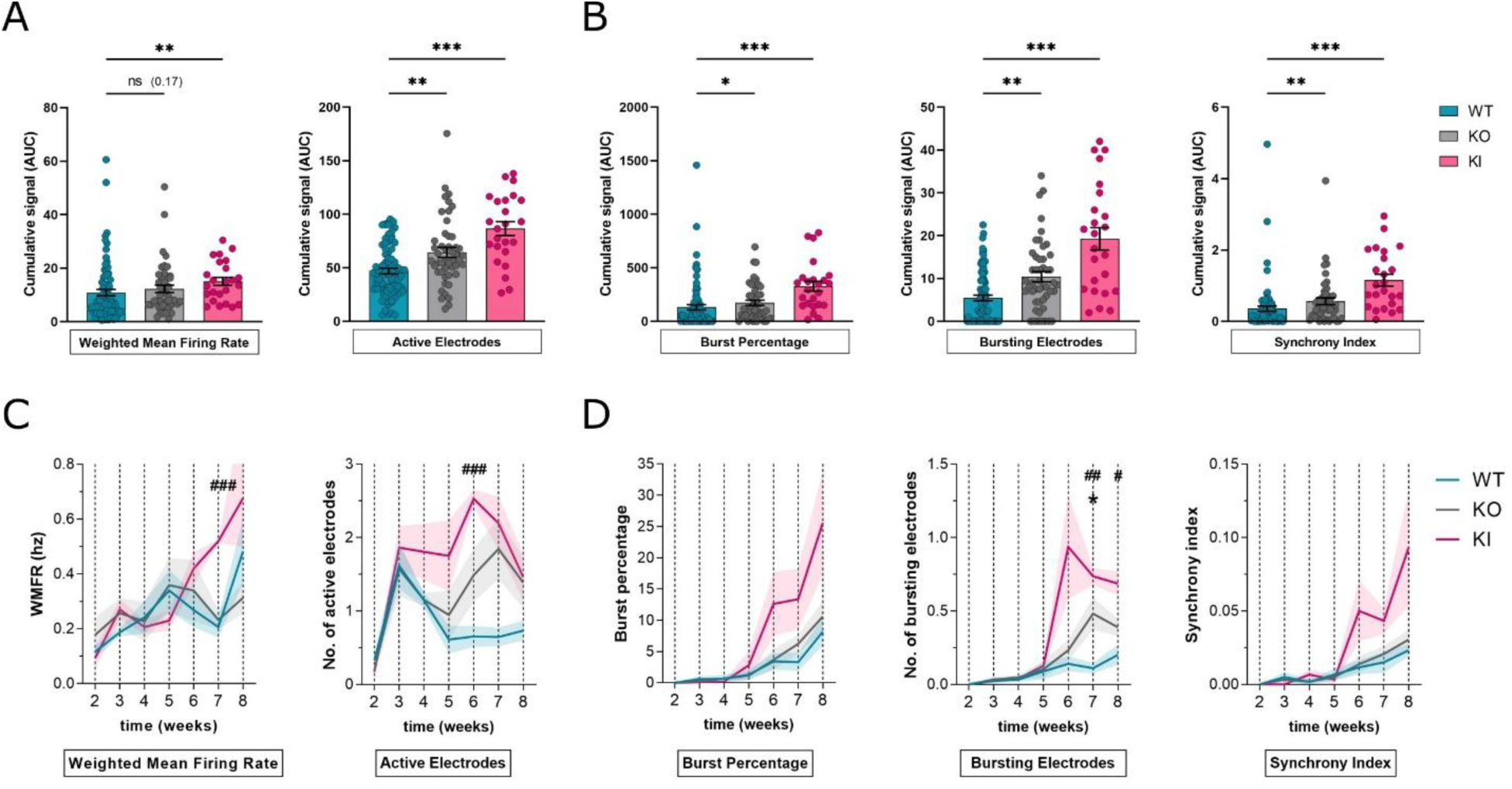
Electrophysiological analysis reveals increased neuronal excitability in SORLA KO and SORLA p.Y1816C mutant neurons; **(A)** Quantification of individual activity parameters recorded by MEA: weighted mean firing rate and number of active electrodes; **(B)** Quantification of network-level parameters: burst percentage, number of bursting electrodes, and synchrony index; **(C)** Time-course visualization of weighted mean firing rate and number of active electrodes over several weeks of neuronal differentiation; **(D)** Time-course analysis of burst percentage, bursting electrodes, and synchrony index across the same period. Each data point represents one well of neuronal culture derived from a different iPSC line. 48, 48, and 24 datapoints are shown from WT, KO, and KI neurons, respectively. Data represent mean ± SEM; Statistical significance was determined using Kruskal–Wallis followed by Dunn’s multiple comparison test or Two-way ANOVA followed by Dunnett’s post-test; *p < 0.05, **p < 0.01, ***p < 0.001. WT vs KO = *; WT vs KI = #. A list of all experimental samples, including iPSC clones used for individual analyses and the number of biological replicates, is provided in Table S2.

Lastly, to explore when these functional changes emerged, we visualized the time course of neuronal maturation on MEA over several weeks. Across multiple measurements, most electrophysiological differences became visible, and often statistically significant, around six weeks of differentiation, highlighting a delayed onset of network dysfunction. The longitudinal evolution of each parameter is presented in **Figures 7C and 7D**. Taken together, these results strongly indicate that both SORLA loss and the AD-causing mutation in *SORL1* p.Y1816C lead to increased neuronal excitability and altered network dynamics in human neurons.

## 4. Discussion

In this study, we systematically investigate the pathogenic impact of the Alzheimer’s-associated *SORL1* p.Y1816C variant using two complementary human-relevant neuronal models: 2D neurons and 3D cerebral organoids. First, we showed that the p.Y1816C variant disrupts SORLA maturation and reduces its proteolytic shedding, without affecting neuronal or organoid differentiation. Subsequently, we demonstrated that both the SORLA KO and KI models exhibit pronounced enlargement of early endosomes in neuronal somas and 3D organoids, confirmed by both Immunofluorescence and ultrastructural TEM analyses. Importantly, this endosomal phenotype was accompanied by an increased accumulation of APP within early endosomes, as well as elevated production of Aβ40 and Aβ42 peptides and deposition of Aβ in both model systems. Moreover, we observed significant neuritic swellings and striking alterations in Rab5+ endosomal transport along axons, including increased stalling and reduced velocity of early endosomes. Importantly, we also showed that APP axonal trafficking is similarly affected, with increased stationary particles and altered velocity in both KO and KI neurons. Lastly, we revealed that SORLA dysfunction leads to deregulated neuronal networks, with increased firing rates, burst activity, and synchrony, most prominently in neurons carrying the p.Y1816C mutation. Together, these results provide mechanistic insight into how SORLA deficiency or mutation disrupts endosomal trafficking and APP processing, ultimately contributing to neuronal dysfunction relevant to AD

## Altered biology of SORLA

At the beginning of our study, we conducted a series of experiments to investigate key aspects of SORLA protein biology, specifically its maturation and shedding, two post-translational processes crucial for the function of SORLA in protein trafficking [33]. Notably, previous studies by us and others have demonstrated that these processes are frequently disrupted in the context of pathogenic *SORL1* variants associated with AD [12,13,17,40]. First, SORLA maturation, which reflects its processing through the Golgi apparatus into a functional, glycosylated form, has been reported to be compromised in several pathogenic variants. In the study by Fazeli et al., we demonstrated that the *SORL1* p.R953C variant exhibited severely impaired maturation, leading to protein retention in the endoplasmic reticulum and loss of downstream trafficking functions [43]. Additional studies have supported the view that reduced SORLA maturation disrupts its ability to interact with the retromer complex and affects APP sorting, thereby contributing to increased amyloidogenic processing [7,18,44]. Importantly, once trafficked to the cell surface, SORLA undergoes proteolytic cleavage by ADAM family proteases, releasing its soluble ectodomain (sSORLA), an indicator of proper surface trafficking. We and others have previously demonstrated that sSORLA shedding is reduced when the receptor is dysfunctional or retained in intracellular compartments [17]. Notably, sSORLA levels were also found to be decreased in cerebrospinal fluid of *SORL1* mutation carriers (de Waal *et al.*, manuscript submitted). Our current findings now expand this understanding by showing that the *SORL1* p.Y1816C variant leads to both impaired maturation and reduced shedding of SORLA in human-relevant models with endogenous *SORL1* expression, providing further functional evidence that this missense variant is pathogenic.

## Endosomal enlargement and APP processing

Given the central role of SORLA in endosomal protein sorting, we next examined how its dysfunction impacts endosomal architecture and APP metabolism. We first focused on endosomal enlargement, a cellular pathology strongly associated not only with *SORL1* mutations but also increasingly recognized as a general upstream mechanism in AD pathogenesis [1]. This theory proposes that enlarged early endosomes represent one of the earliest detectable abnormalities in AD, preceding amyloid plaque deposition and Tau pathology. Disruption of endosomal function may impair trafficking of key neuronal cargo, including APP, neurotrophic receptors, and lipids, thereby triggering downstream neurodegenerative cascades [3,6,34]. Indeed, endosomal enlargement has been extensively reported in models of *SORL1* deficiency or mutants [12,13,15,43], as well as in the context of other AD-related genes such as *PSEN1*, *APP*, and *APOE4* [14,34,45]. Notably, this phenotype is especially relevant to APP processing, as β-secretase (BACE1), the enzyme initiating the amyloidogenic cleavage pathway, predominantly acts within early endosomes [46– 48]. Thus, retention of APP in enlarged endosomes increases its accessibility to BACE1, promoting the generation and secretion of Aβ40 and Aβ42 peptides, a central feature of AD pathology. Importantly, previous studies using *SORL1* KO models, overexpression systems, and iPSC-derived neurons have confirmed that SORLA loss results in increased APP accumulation in endosomes and elevated Aβ secretion [7,10,12]. Consistent with these findings, our data now show that the *SORL1* p.Y1816C mutation induces significant enlargement of early endosomes, increased APP retention within these compartments, and enhanced production of Aβ40 and Aβ42 peptides. Notably, building on previous work, we provide the first ultrastructural evidence of early endosome enlargement in human neurons using TEM, offering direct visual confirmation of altered endosomal architecture. Moreover, we demonstrate that these phenotypes are not confined to 2D neuronal cultures but are faithfully recapitulated in 3D cerebral organoids. This represents a key advancement, as it confirms that SORLA-related endosomal dysfunction is preserved in more complex, tissue-like models that better mimic the cellular architecture and network environment of the human brain. Thus, the replication of these defects in cerebral organoids highlights the robustness of the observed phenotypes and also underscores the utility of 3D systems for modeling early-stage AD mechanisms.

## Compromised axonal biology and axonal transport in AD

While somatic vesicle trafficking has been the primary focus of many studies on SORLA function, far less is known about its potential roles in the axonal compartment. Indeed, neurons rely on precisely regulated long-range intracellular transport to move organelles, receptors, and signaling molecules between the cell body and distant axonal or dendritic compartments. These processes depend on microtubule-based transport systems powered by motor proteins, dyneins and kinesins [21]. Importantly, disruption of these mechanisms has been increasingly recognized as an early and critical feature of neurodegenerative diseases, including AD. Interestingly, one well-documented hallmark of compromised axonal integrity is the formation of neuritic or axonal swellings, focal enlargements along the axon associated with vesicular accumulation, impaired transport, and neurodegeneration [2,23,41]. Moreover, such swellings have been observed across a range of neurodegenerative diseases, including AD (reviewed in [22]). Interestingly, among the cargoes trafficked along axons, Rab5-positive early endosomes play a significant role, carrying molecules such as APP, TrkB, and recycling receptors in both anterograde and retrograde directions [49,50]. Disruption of this transport system has been shown in AD models carrying the APP Swedish mutation, where endosomes and APP particles exhibit abnormal stalling and impaired motility [20]. Our current findings now extend this knowledge by revealing that both *SORL1* p.Y1816C mutant and SORLA knock-out neurons exhibit clear morphological and functional signs of axonal dysfunction. We observed pronounced neuritic swellings, often containing enlarged endosomes, an indicator of disturbed axonal trafficking. Live-cell imaging confirmed that Rab5+ endosomes in SORLA-deficient neurons exhibit abnormal clustering, increased stalling, and reduced bidirectional transport velocities. Parallel analyses of APP trafficking revealed similar impairments, including an increased proportion of stationary APP-GFP particles. These deficits in axonal transport were accompanied by a significant increase in the number, width, and length of neuritic swellings, further supporting a direct link between SORLA dysfunction and axonal pathology. Taken together, our results demonstrate that SORLA is essential not only for somatic vesicle trafficking but also for maintaining axonal homeostasis and transport dynamics, thereby implicating SORL1 dysfunction as a mechanistic driver of axonal pathology in AD.

## Hyperexcitability and AD

Finally, we examined neuronal hyperexcitability, a functional phenotype that emerges early in AD and may contribute to disease progression. Multiple studies have demonstrated that neurons carrying AD-causing mutations, including variants in *APP, PSEN1*, and *PSEN2*, exhibit aberrant network activity and increased excitability. For instance, studies on human iPSC-derived neurons and organoids with familial AD mutations have reported increased spontaneous firing and calcium oscillation rates [14,24]. Similarly, mouse models of AD, such as 5xFAD and APP/PS1 mice, show spontaneous epileptiform activity, suggesting that neuronal hyperexcitability precedes overt neurodegeneration [25,26]. Clinical observations also support this, as increased subclinical seizure activity and cortical hyperexcitability have been detected in individuals with early-onset or autosomal dominant AD, even in the preclinical stages [51,52]. Notably, a study by Zhou et al. reported that iPSC-derived neurons carrying the APP Swedish mutation display increased spontaneous synaptic activity, which they attributed to the action of Aβ peptides accumulating in the culture media, acting in a paracrine manner to modulate synaptic excitability [27]. However, despite SORLA being known to regulate APP trafficking and Aβ generation, its role in modulating the electrophysiological properties of neurons has remained largely unexplored. Only one prior study by the Young lab reported increased excitability in SORL1 knock-out human neurons [15]. Our data now demonstrate that both *SORL1* p.Y1816C and SORLA knock-out neurons exhibit deregulated electrophysiology. This included elevated firing rates, a greater number of active and bursting electrodes, and increased synchrony across the network.

Importantly, these findings are further corroborated by Williams *et al.* (submitted back-to-back with this manuscript), who not only observed hyperexcitability in *SORL1*-deficient neurons, confirming our findings, but also describe that this hyperexcitability is not solely due to the overproduction of Amyloid-β but rather due to the mis-trafficking of synaptic proteins. Together, our findings and those of Williams *et al.* add compelling evidence that *SORL1* dysfunction disrupts multiple layers of neuronal biology, ranging from subcellular trafficking to synaptic function, reinforcing the emerging view that endosomal recycling defects are central, upstream drivers in Alzheimer’s disease pathogenesis.

## Conclusion

In summary, our study demonstrates that the Alzheimer’s-linked *SORL1* p.Y1816C variant impairs multiple facets of neuronal function, including SORLA maturation, endosomal trafficking, APP processing, axonal transport, and neuronal excitability. Using both 2D and 3D human neuronal models, we show that SORLA dysfunction leads to hallmark AD-related cellular phenotypes such as early endosome enlargement, Aβ accumulation, axonal swellings, and network hyperexcitability. These findings underscore the central role of SORL1 in maintaining neuronal homeostasis and support the notion that endosomal trafficking defects are a key pathogenic mechanism in AD.

## List of abbreviations

AD: Alzheimer’s Disease
Aβ: Amyloid-β
APP: Amyloid Precursor Protein
BACE1: Beta-Site APP Cleaving Enzyme 1
CNM: Cortical Neuron Medium
CSF: Cerebrospinal Fluid
ED: Electron Dense Endosomes
EL: Electron Lucent Endosomes
ER: Endoplasmic Reticulum
Ipsc: Induced Pluripotent Stem Cell
IM: Induction Medium
KO: Knock-Out
KI: Knock-In
MEA: Multielectrode Array
PSEN: Presenilin
SEM: Scanning Electron Microscopy
Ssorla: Soluble SORLA ectodomain
SORL1: Sortilin-Related Receptor 1 (gene)
SORLA: Sortilin-Related Receptor with A-type Repeats (protein encoded by SORL1)
TEM: Transmission Electron Microscopy
3Fn: Fibronectin type III domain

## Acknowledgements

The authors would like to thank Prof. Ales Hampl, Prof. Henne Holstege, and Dr. Sven van der Lee for their continuous support. We acknowledge the CF Genomics supported by the NCMG research infrastructure (LM2023067 funded by MEYS CR) for their support with flow cytometry and CELLIM core facility, supported by the Czech-BioImaging large research infrastructure project (LM2023050 funded by MEYS CR), for their support in acquiring imaging data used in this work.Veronika Pospisilova and Tereza Vanova acknowledge the support from Alzheimer NF, Sona Cesnarikova acknowledges the support by a PhD Talent Fellowship.

---

**Supplementary Figure S1:**
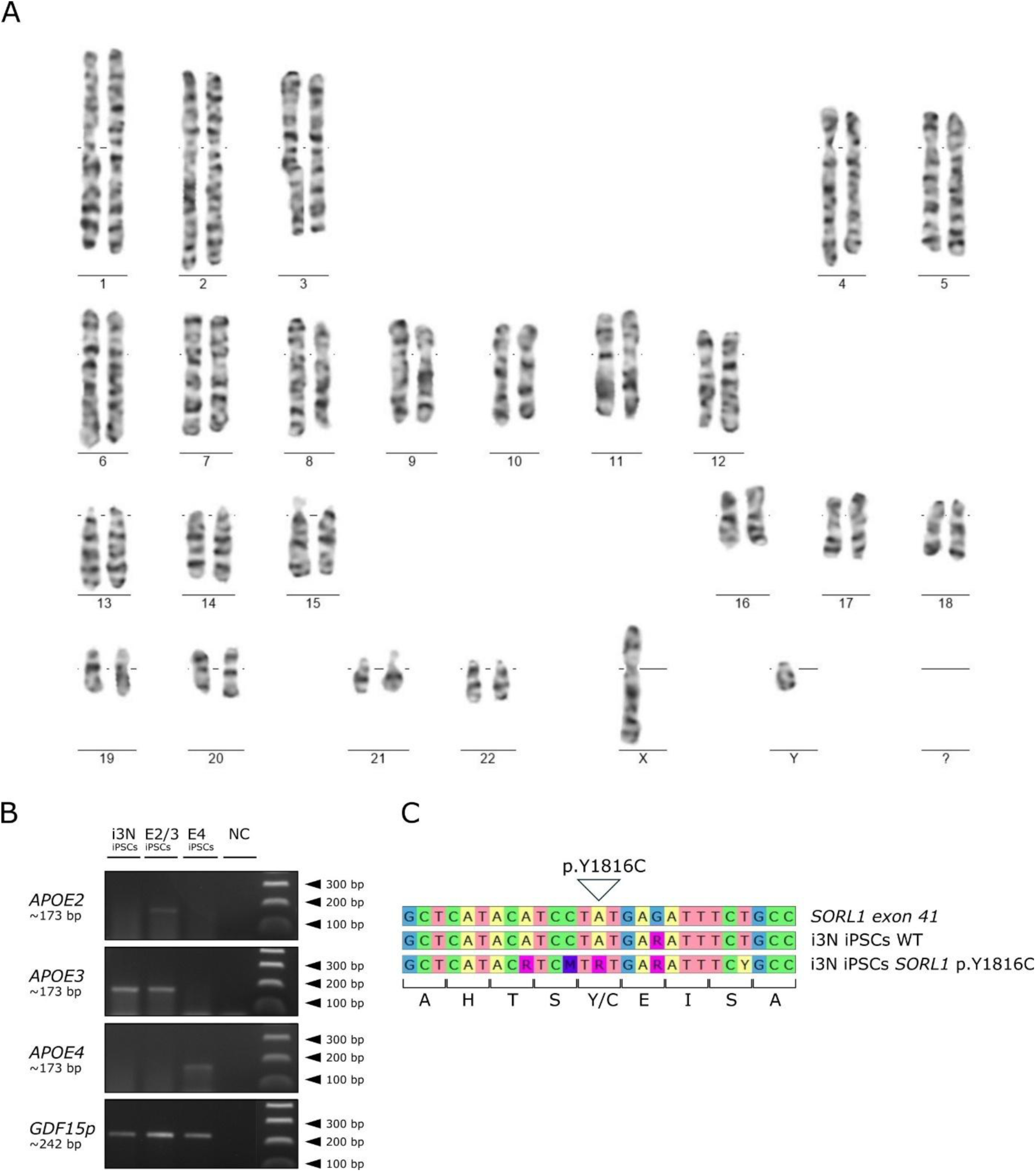
Genetic validation and characterization of *SORL1* gene-edited iPSC lines. **(A)** Representative karyotype analysis showing a normal chromosomal profile across all gene-edited lines. **(B)** APOE genotyping shows a homozygous APOE3/3 genotype in i3N iPSC line **(C)** Sanger sequencing showing introduction of the heterozygous *SORL1* p.Y1816C point mutation in the KI line. A list of all experimental samples, including iPSC clones used for individual analyses and the number of biological replicates, is provided in **Table S2**

**Supplementary Figure S2.**
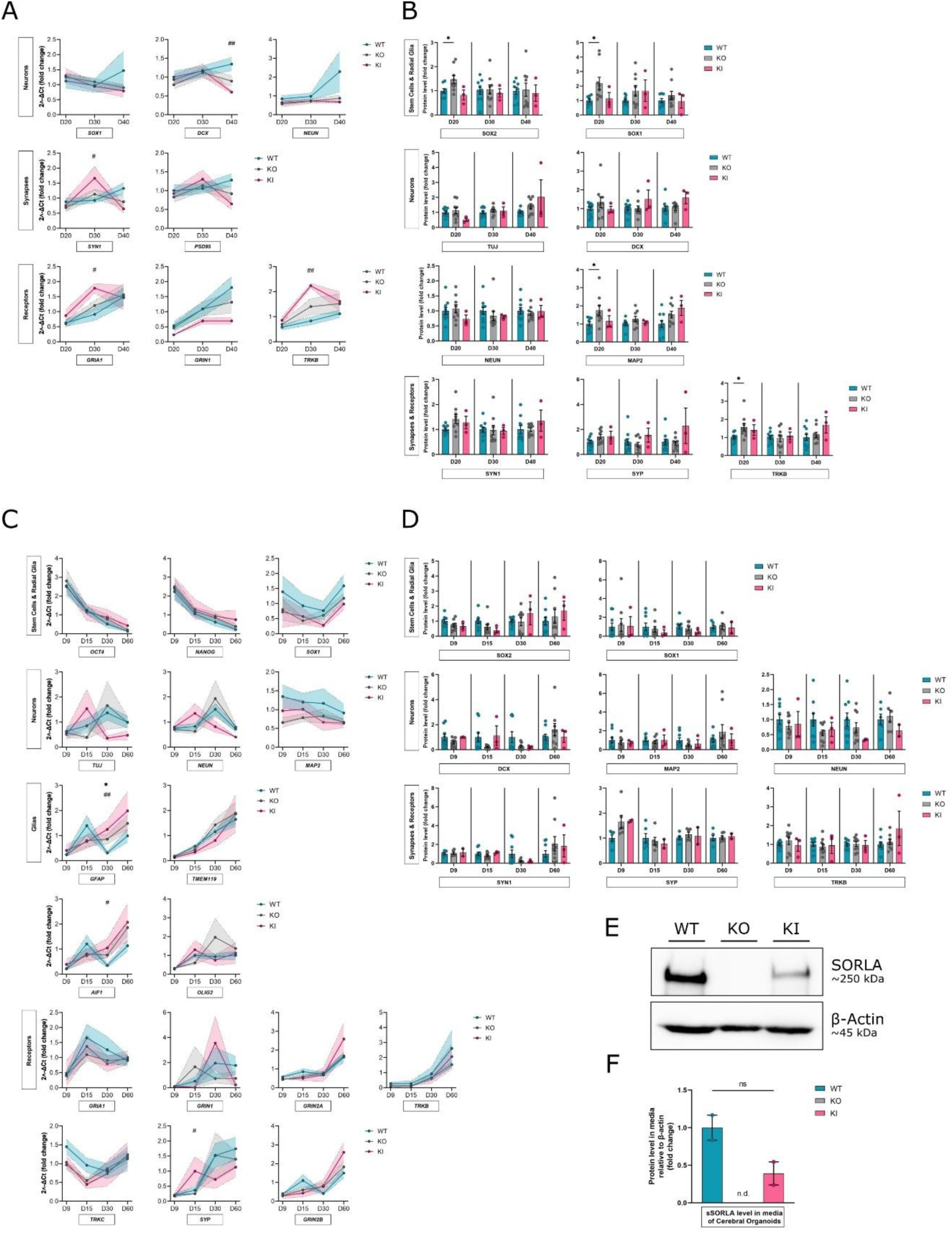
Supporting data for neuronal marker expression, SORLA levels, and sSORLA detection in neurons and organoids. **(A,B)** Additional validation of progenitor and neuronal marker expression at the mRNA and protein levels in 2D neuron cultures across WT, KO, and KI lines. **(C, D)** Gene and protein expression analysis of pluripotency, progenitor, neuronal, synaptic, and glial markers in cerebral organoids. (E) Western blot analysis of culture medium from day 60 cerebral organoids of sSORLA in KO and KI-derived organoids compared to WT; (F) Densitometric quantification of sSORLA levels from (E). Each dot in (A) and (C) represents the average 2˄-ΔCt; each dot in (B), (D), and (F) represents one biological replicate (n ≥ 2). Data are presented as mean ± SEM. Statistical significance was assessed using one way ANOVA followed by Dunnett’s Tukey’s multiple comparison test, Kruskal–Wallis followed by Dunn’s multiple comparison test or t-test; *p < 0.05, **p < 0.01, WT vs KO = * WT vs KI = #. A list of all experimental samples, including iPSC clones used for individual analyses and the number of biological replicates, is provided in Table S2.

**Supplementary Figure S3.**
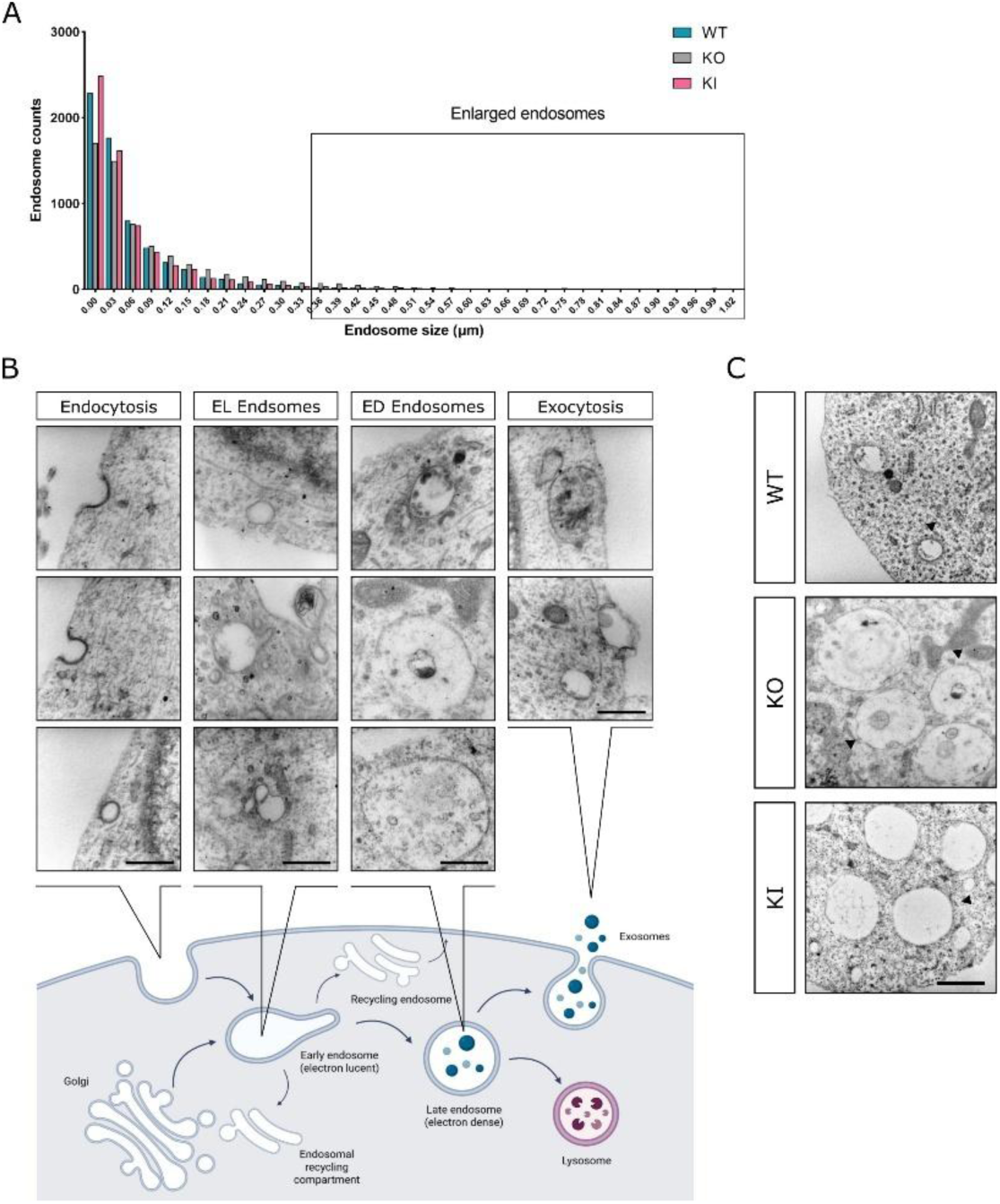
Analyses of endosomal morphology and spatial distribution across genotypes. **(A)** Size distribution of all detected EEA1-positive endosomes in 2D neurons across WT, KI, and KO lines; **(B)** Classification of endolysosomal structures identified in TEM images, including endocytosis of clathrin-coated pits and vesicles, electron-lucent (EL) endosomes, electron-dense (ED) endosomes, and vesicles undergoing exocytosis; **(C)** Representative TEM fields showing clustering of enlarged endosomes within the neuronal soma in WT, KO, and KI neurons; TEM scale bar: 500 nm (B) 1000 nm (C). A list of all experimental samples, including iPSC clones used for individual analyses and the number of biological replicates, is provided in **Table S2.**

**Figure S4.**
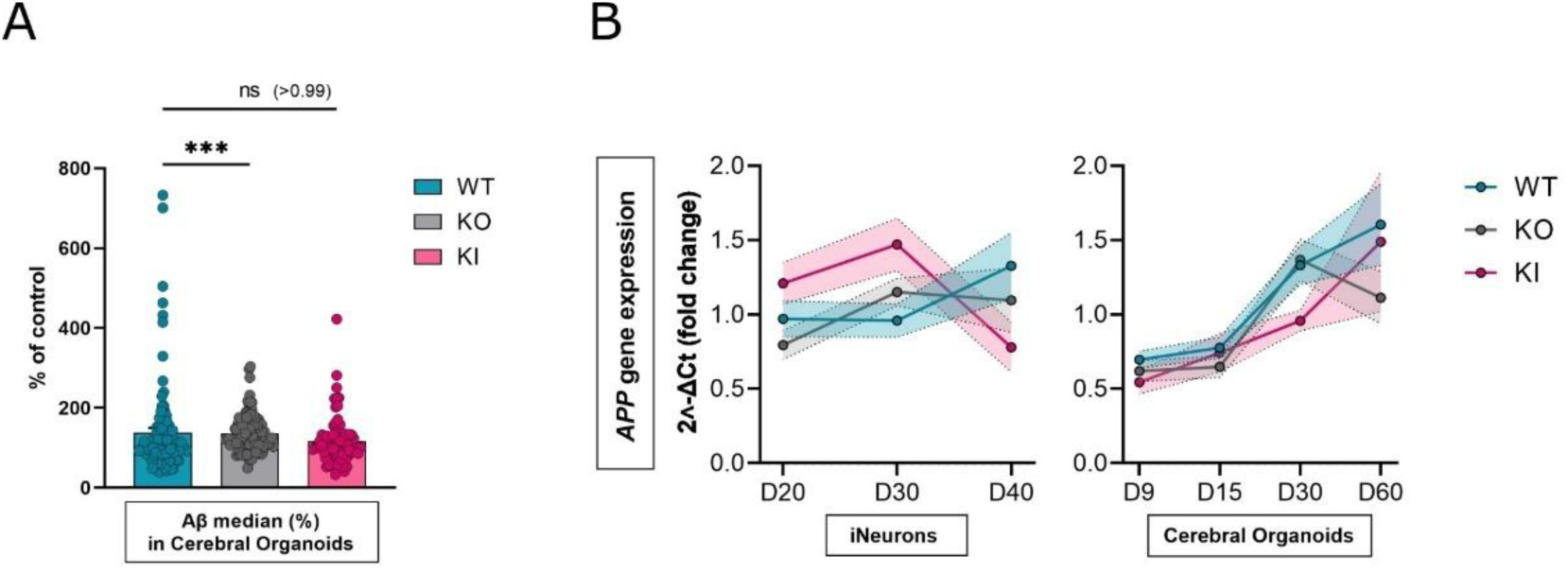
Additional analyses of Aβ cluster size and APP expression in neurons and organoids. **(A)** Median volume of Aβ particles in cerebral organoids across WT, KI, and KO genotypes. The average signal intensity of the control samples was used as a reference for comparing all analyzed data; **(B)** APP mRNA expression in neurons and organoids across WT, KI, and KO lines, as measured by qRT-PCR. Each dot represents one biological replicate (n ≥ 3). Data represent mean ± SEM. Statistical significance was assessed using one-way ANOVA followed by Dunnett’s Tukey’s multiple comparison test, Kruskal–Wallis followed by Dunn’s multiple comparison test or t-test; ***p < 0.001. A list of all experimental samples, including iPSC clones used for individual analyses and the number of biological replicates, is provided in **Table S2**

**Figure S5.**
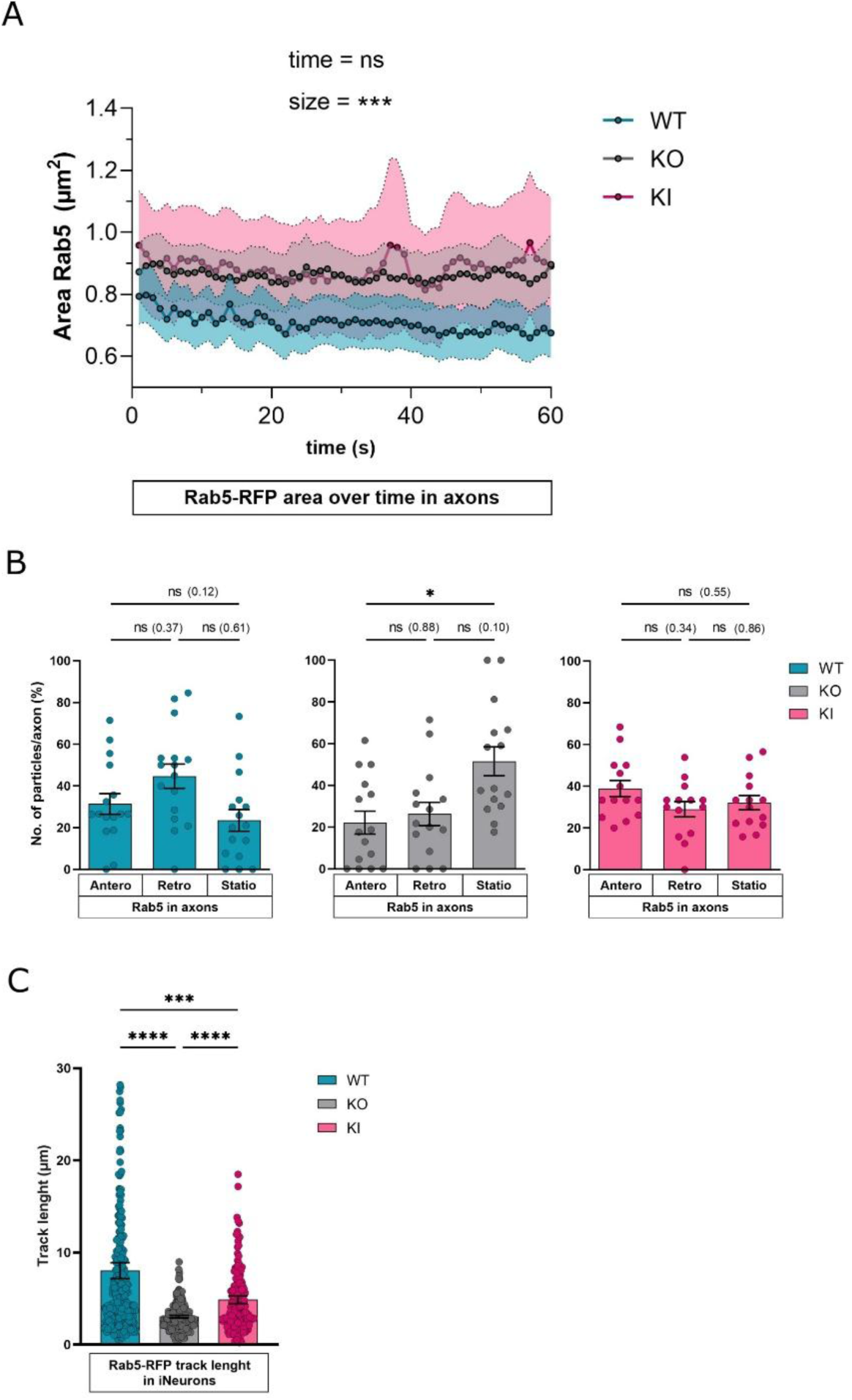
Supplementary quantification of Rab5a-positive endosome dynamics in WT, KI, and KO axons. **(A)** Area of Rab5+ particles over time in WT, KO, and KI; **(B)** Proportion of Rab5+ particles moving anterogradely, retrogradely, or remaining stationary in WT, KI, and KO neurons, quantified per axon; **(C)** Track length of individual Rab5+ endosomes, showing total distance traveled by individual endosomes. Data represent mean ± SEM (in A represented by shaded area). Statistical significance was assessed using one-way ANOVA followed by Dunnett’s Tukey’s multiple comparison test and two-way ANOVA followed by Holm-Šidak’s multiple comparisons test; *p < 0.05, **p < 0.01. A list of all experimental samples, including iPSC clones used for individual analyses and the number of biological replicates, is provided in **Table S2**

**Figure S6.**
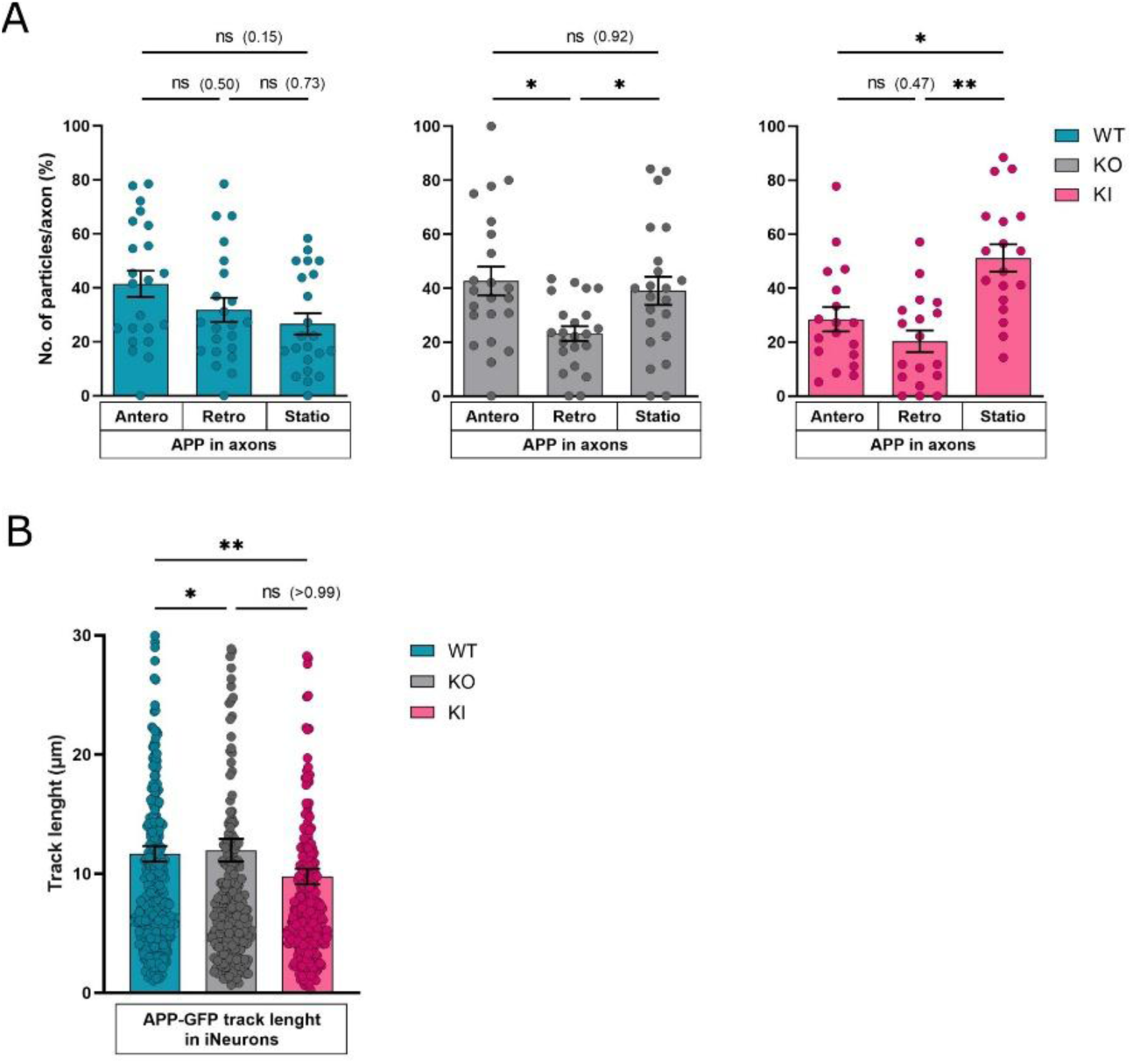
Additional transport metrics of APP-GFP. **(A)** Proportion of APPwt-GFP particles moving anterogradely, retrogradely, or remaining stationary in WT, KI, and KO neurons, quantified per axon. **(B)** Track length of individual GFP signal particles, showing total distance traveled by APP. Each dot represents one axon. A total of 20 axons were measured in three independent biological replicates. Data represent mean ± SEM. Statistical significance was assessed using one-way ANOVA followed by Dunnett’s Tukey’s multiple comparison test; *p < 0.05, **p < 0.01. A list of all experimental samples, including iPSC clones used for individual analyses and the number of biological replicates, is provided in Table S2.

## Supplementary tables

**Table S1:**
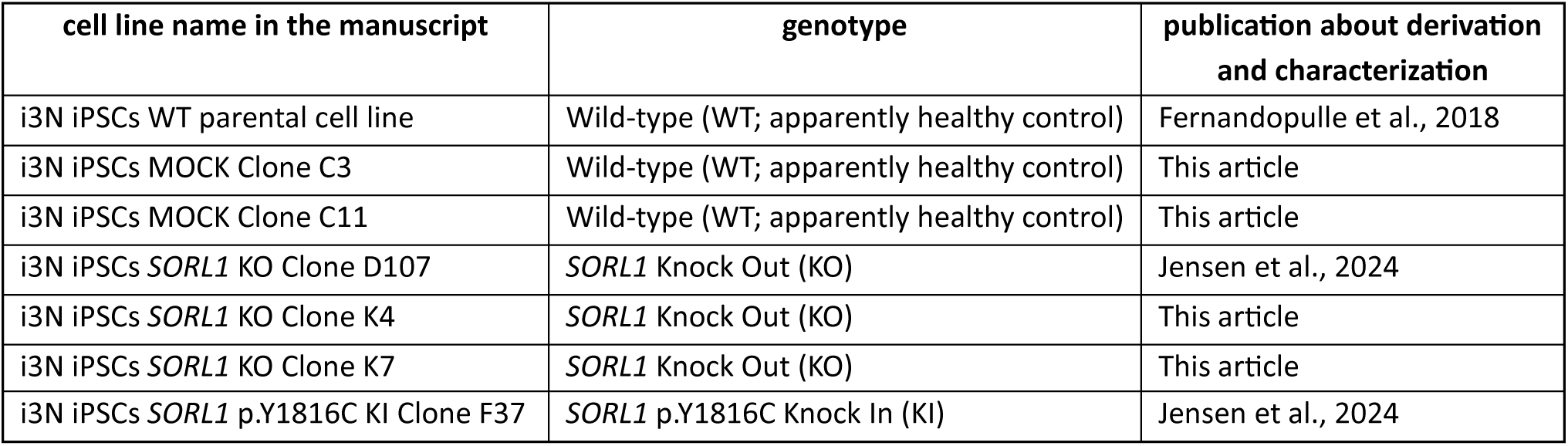
Cell lines used.

| cell line name in the manuscript | genotype | publication about derivation and characterization |
| --- | --- | --- |
| i3N iPSCs WT parental cell line | Wild-type (WT; apparently healthy control) | Fernandopulle et al., 2018 |
| i3N iPSCs MOCK Clone C3 | Wild-type (WT; apparently healthy control) | This article |
| i3N iPSCs MOCK Clone C11 | Wild-type (WT; apparently healthy control) | This article |
| i3N iPSCs *SORL1* KO Clone D107 | *SORL1* Knock Out (KO) | Jensen et al., 2024 |
| i3N iPSCs *SORL1* KO Clone K4 | *SORL1* Knock Out (KO) | This article |
| i3N iPSCs *SORL1* KO Clone K7 | *SORL1* Knock Out (KO) | This article |
| i3N iPSCs *SORL1* p.Y1816C KI Clone F37 | *SORL1* p.Y1816C Knock In (KI) | Jensen et al., 2024 |

**Table S2:**
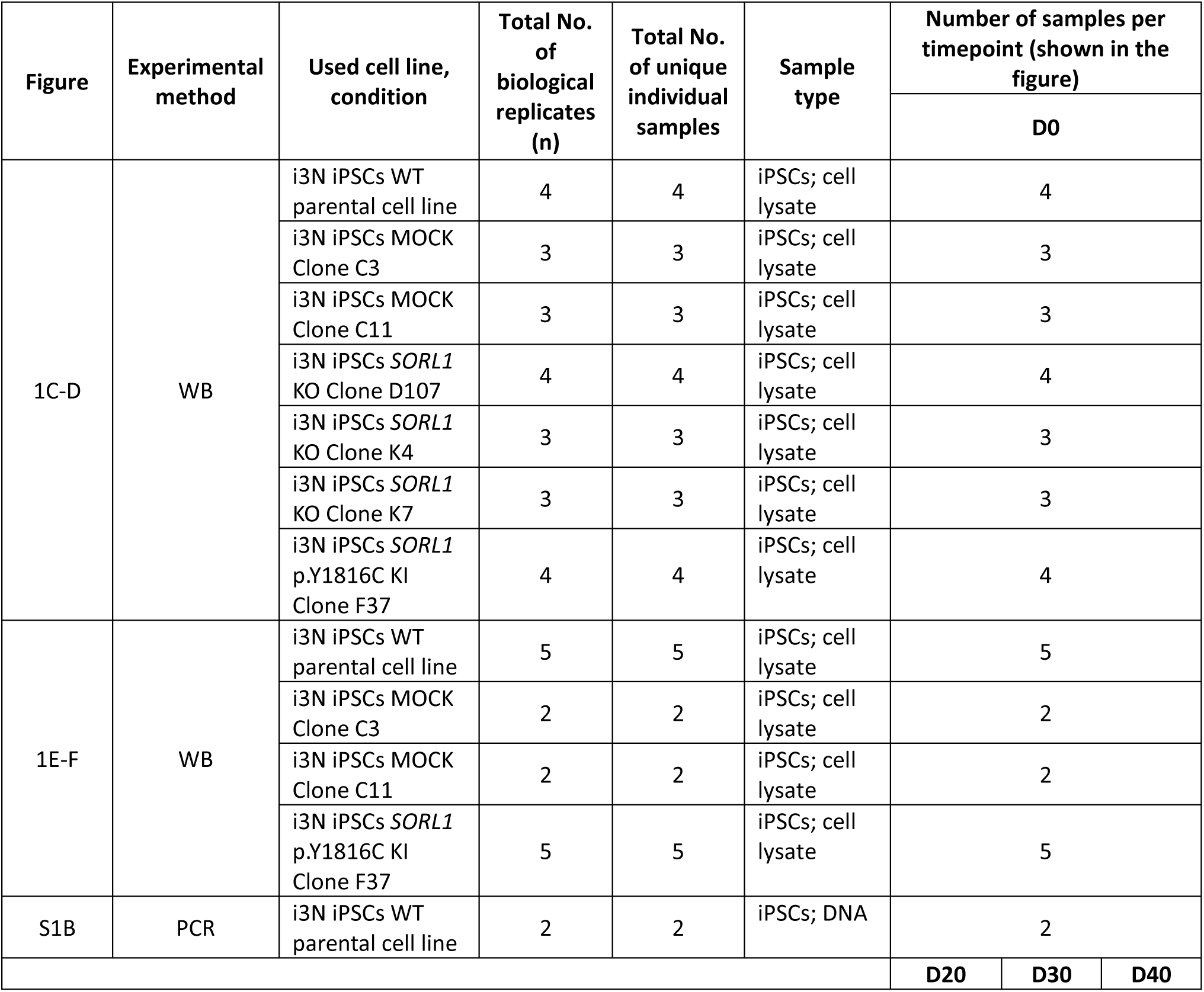
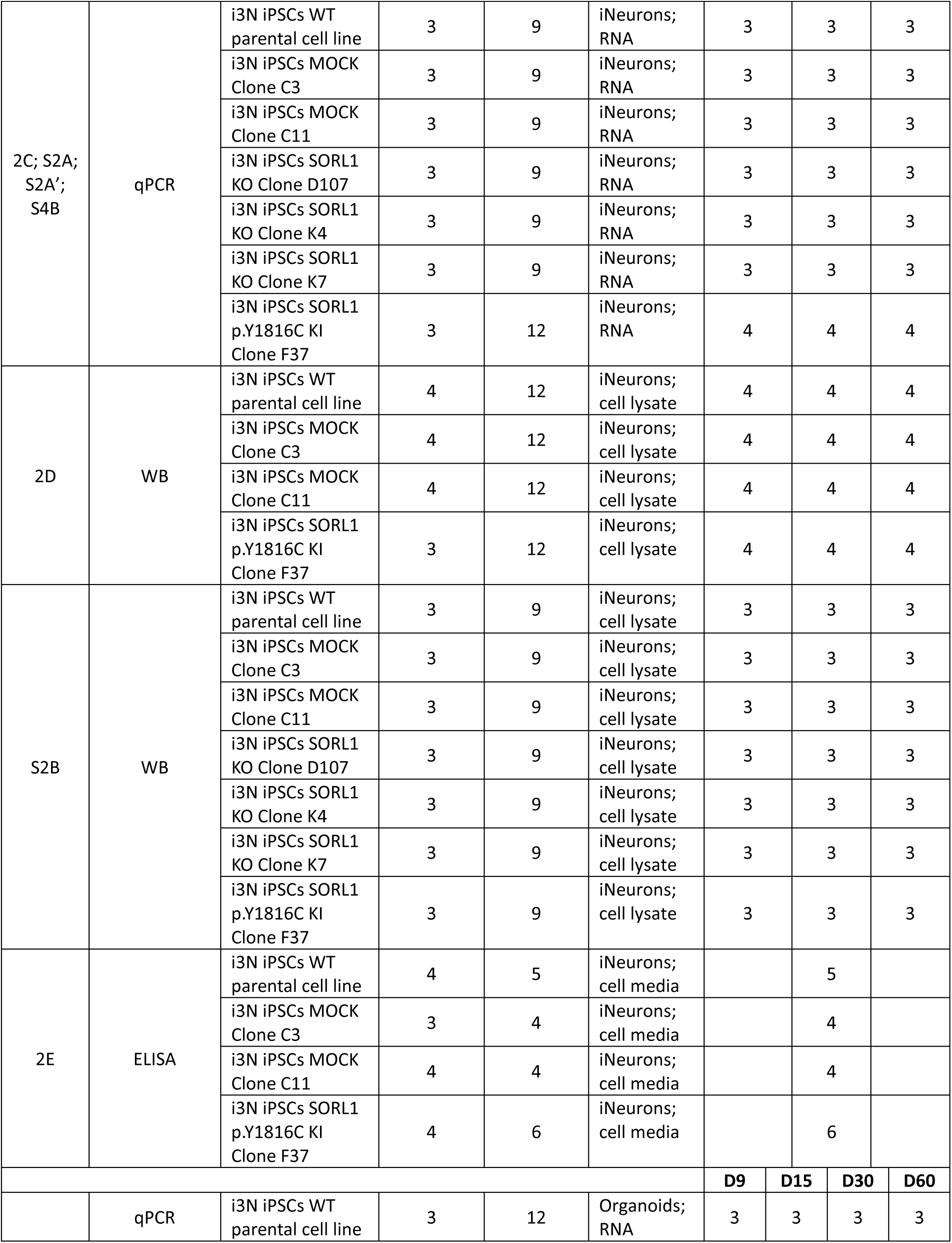
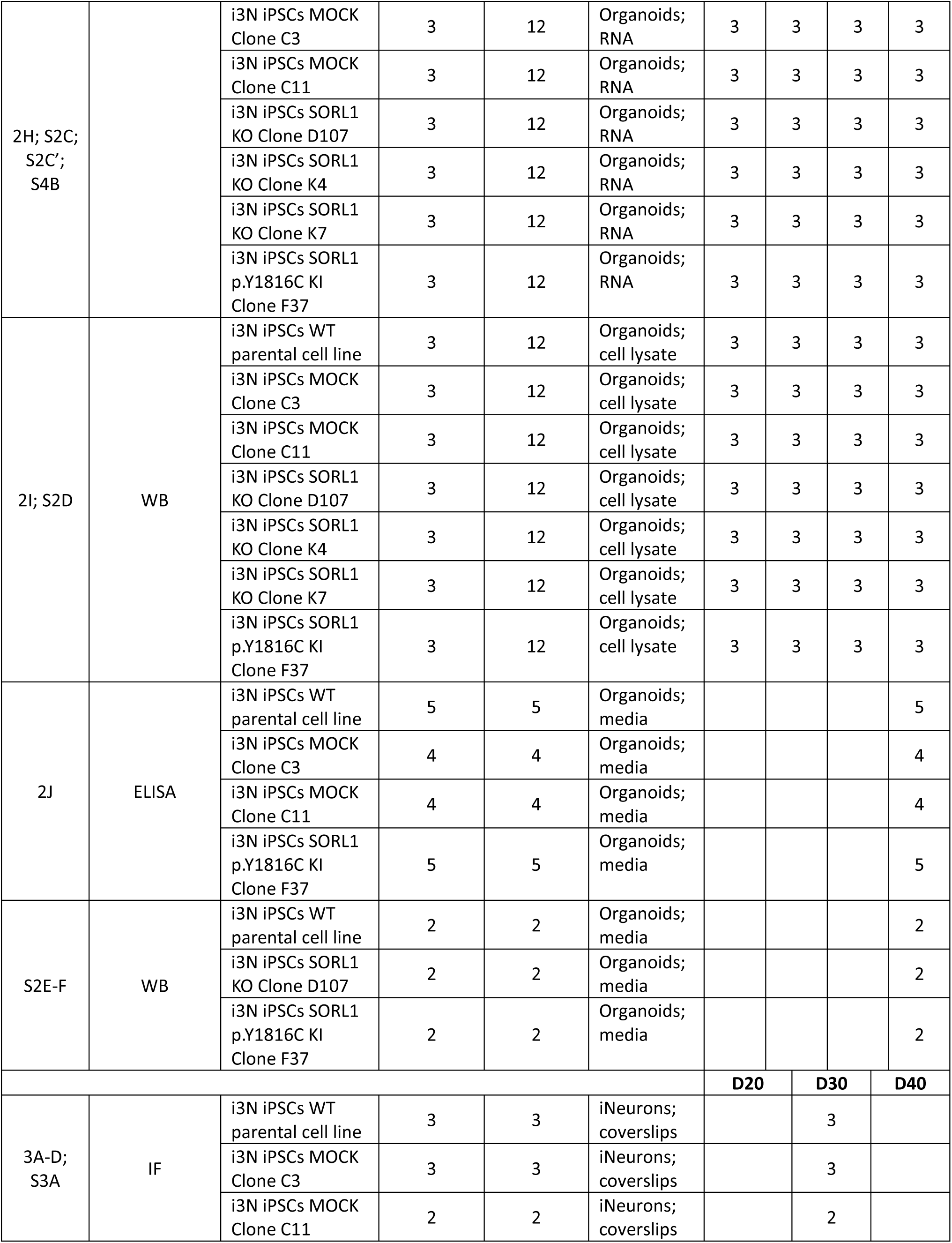
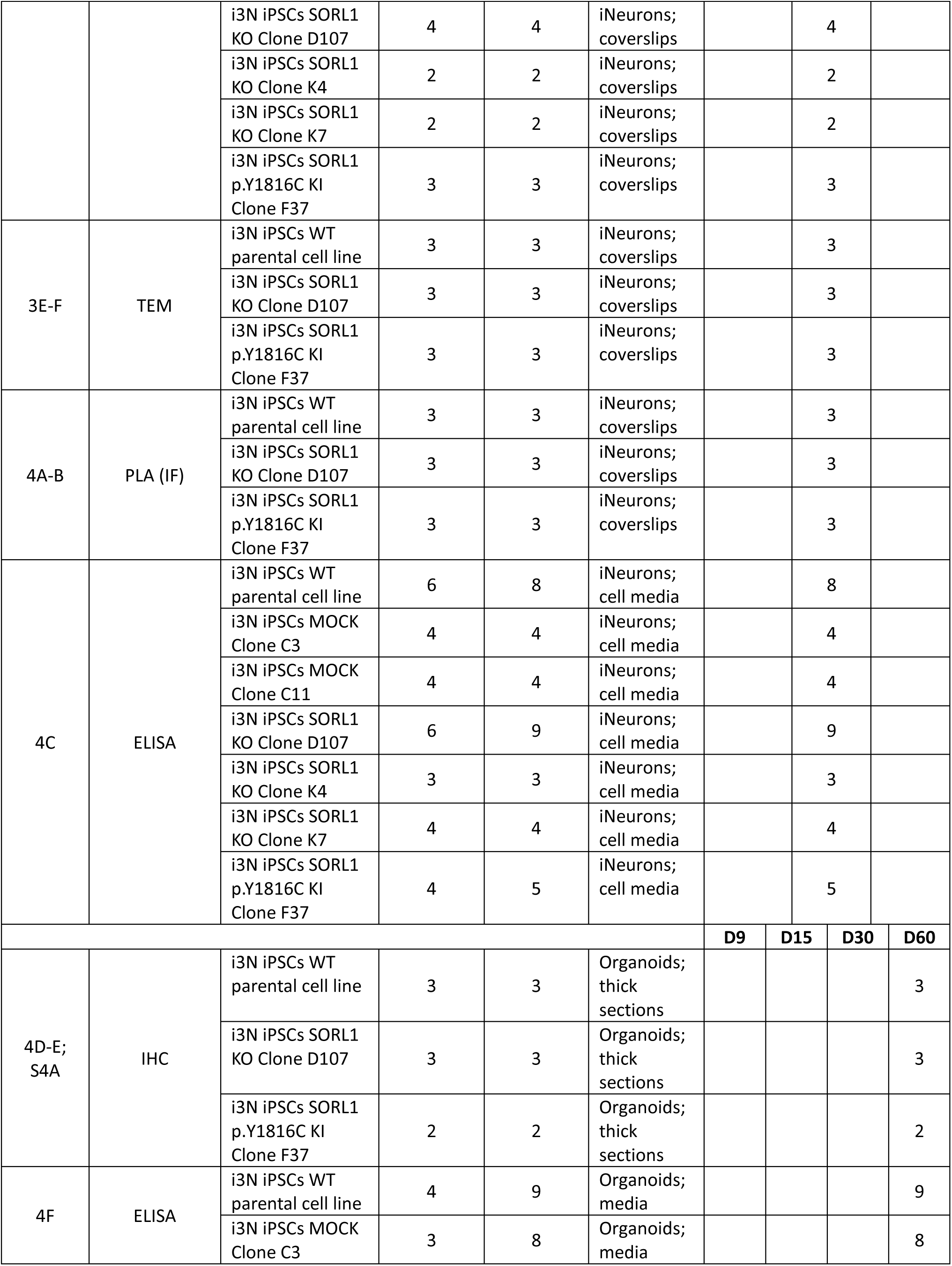
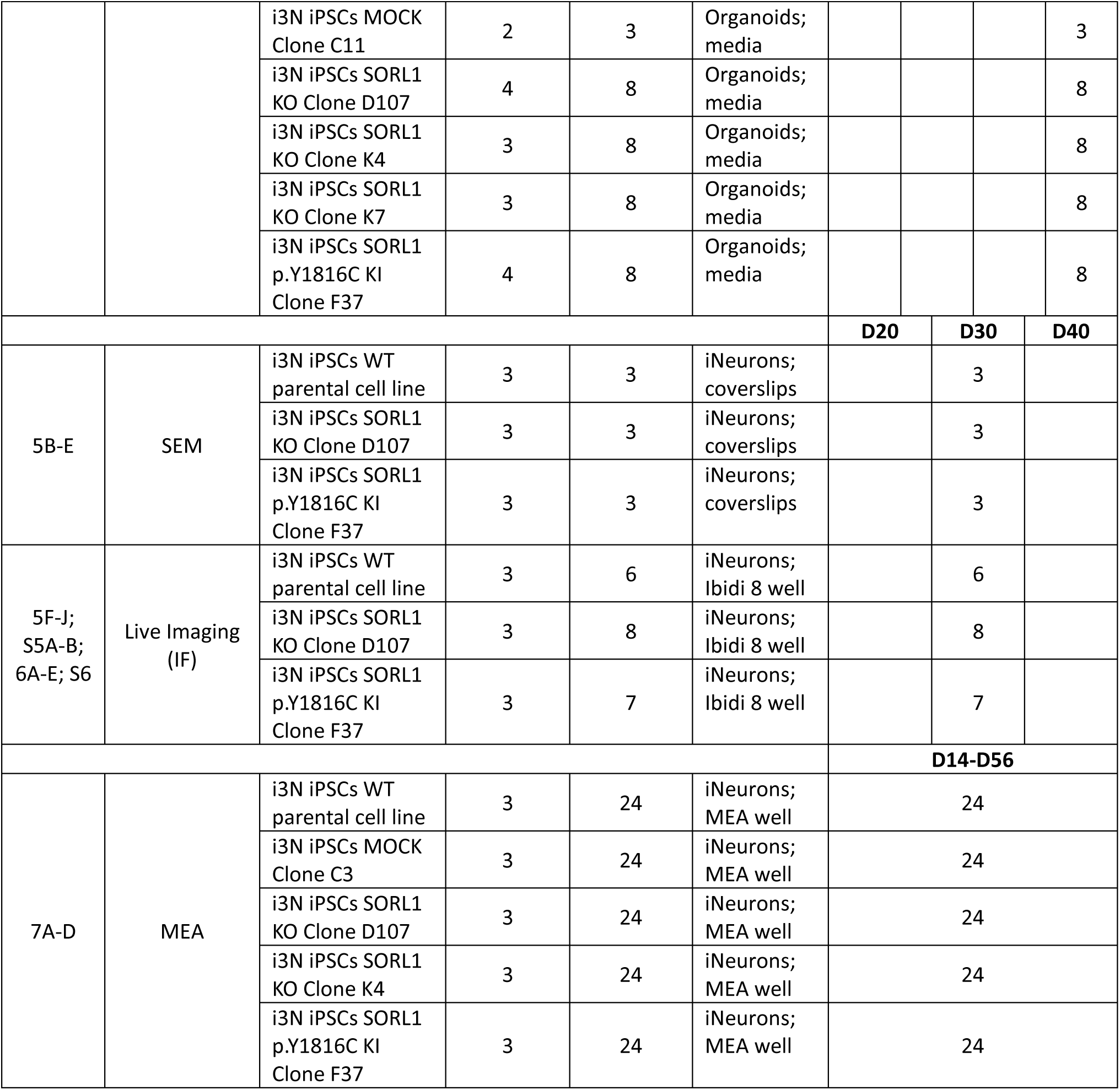
Number of replicates per experiment.

**Table S3:**
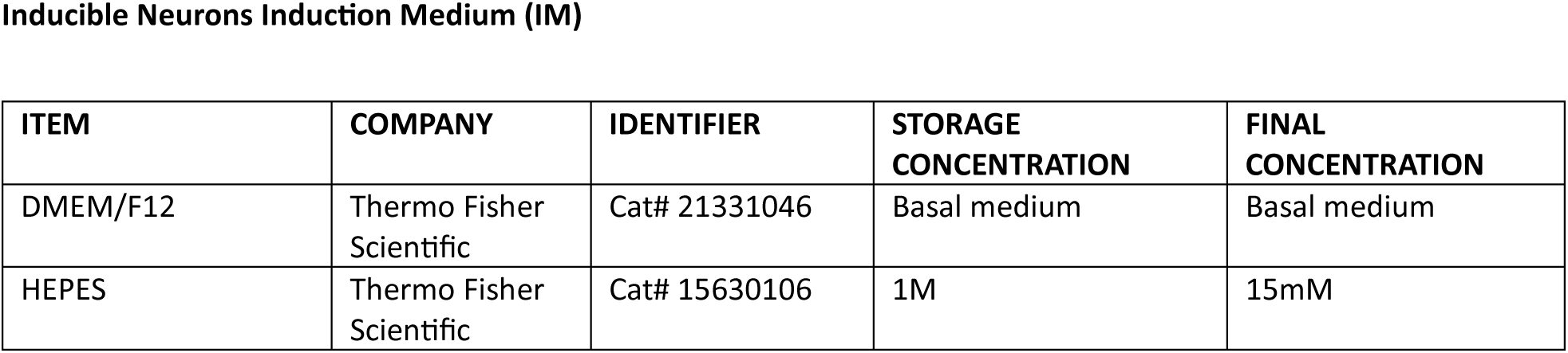
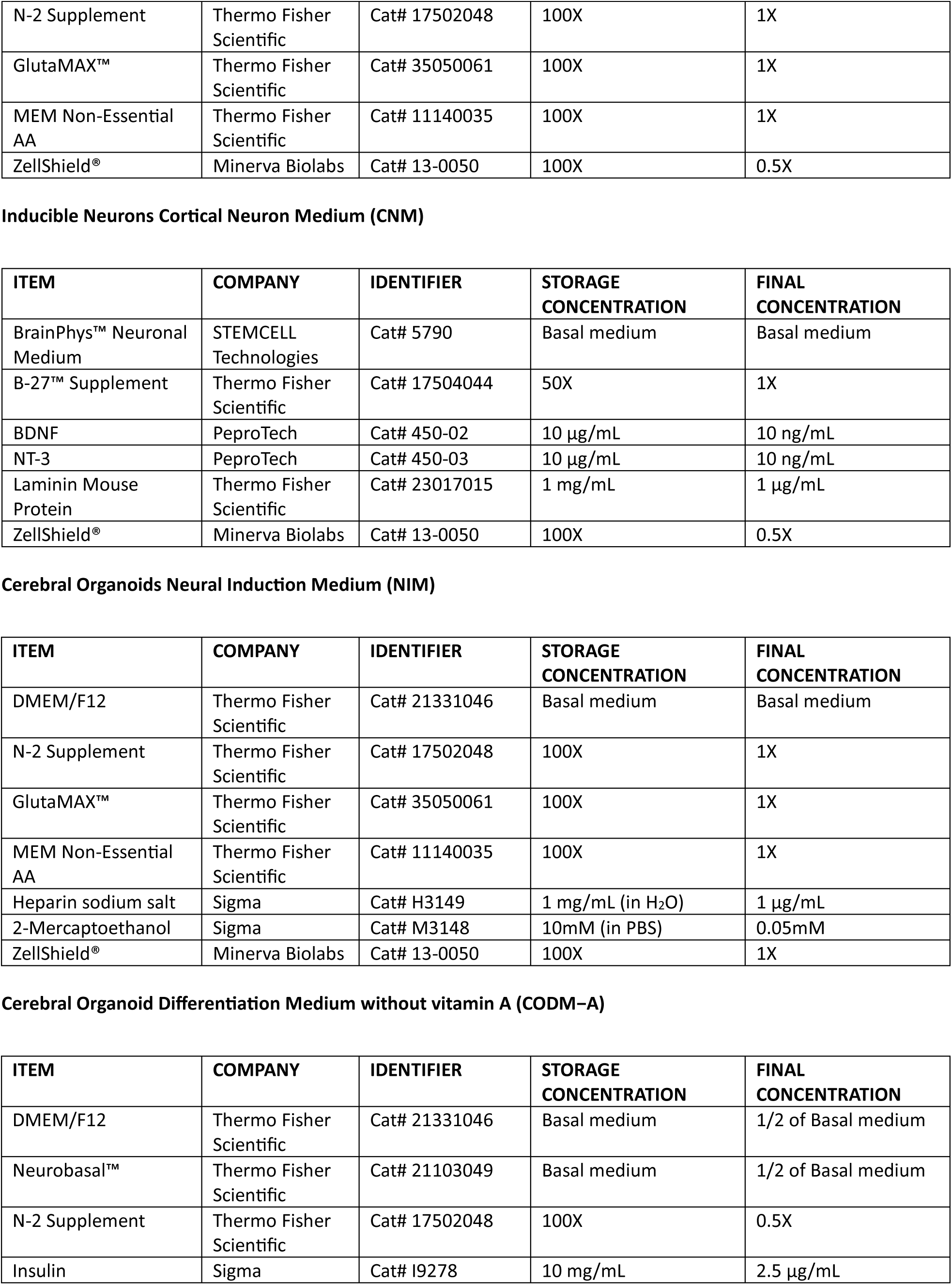
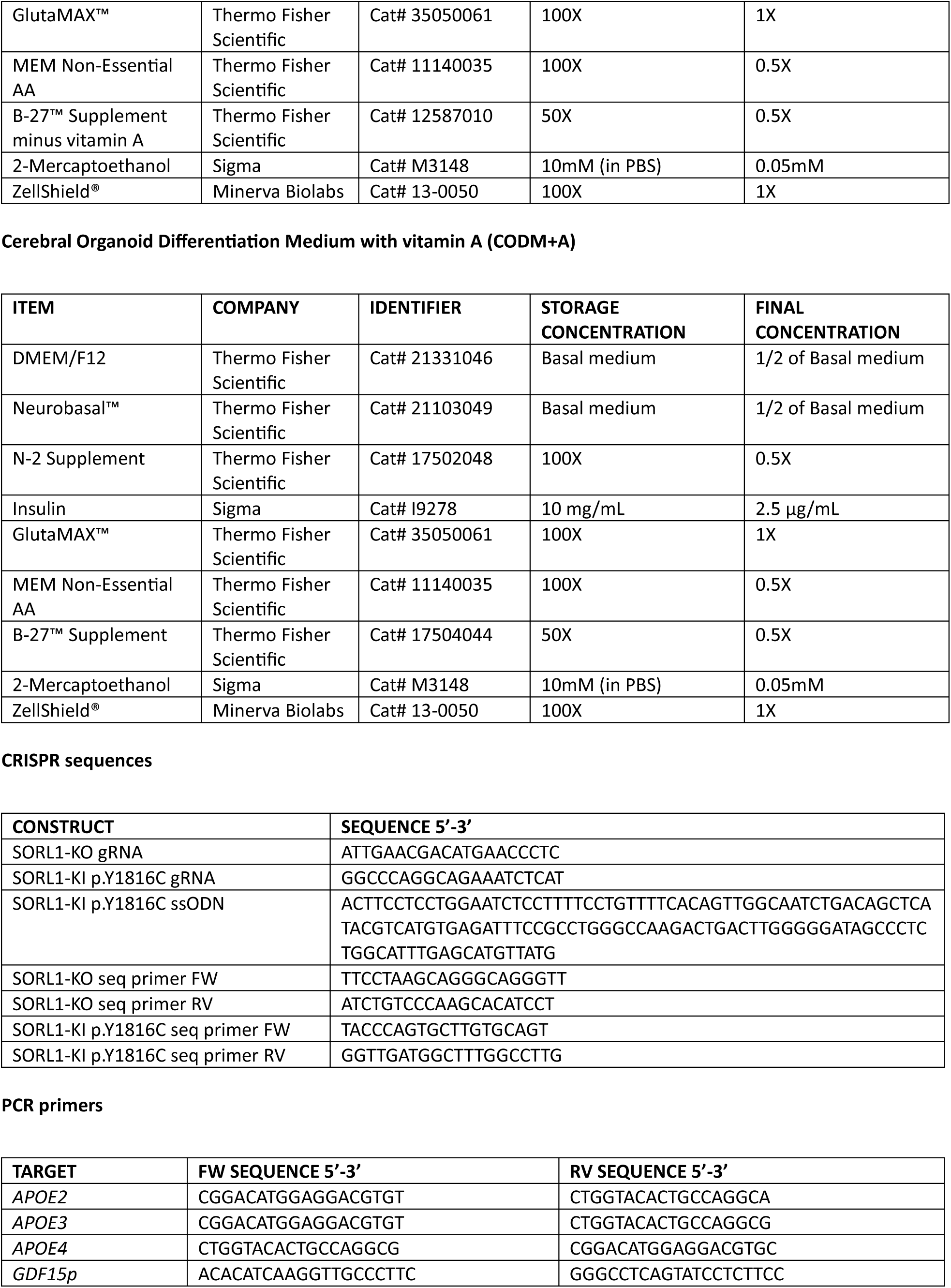
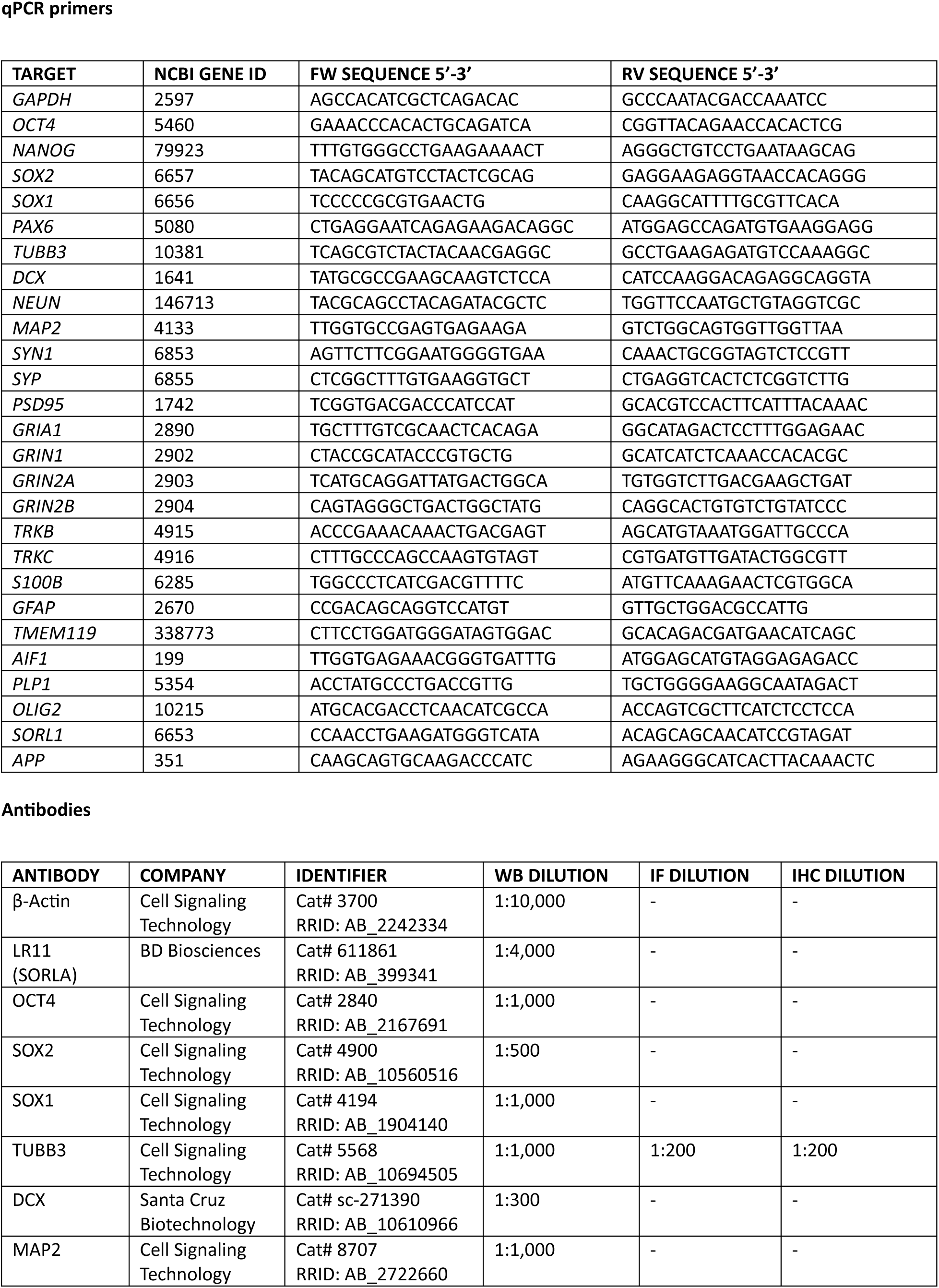
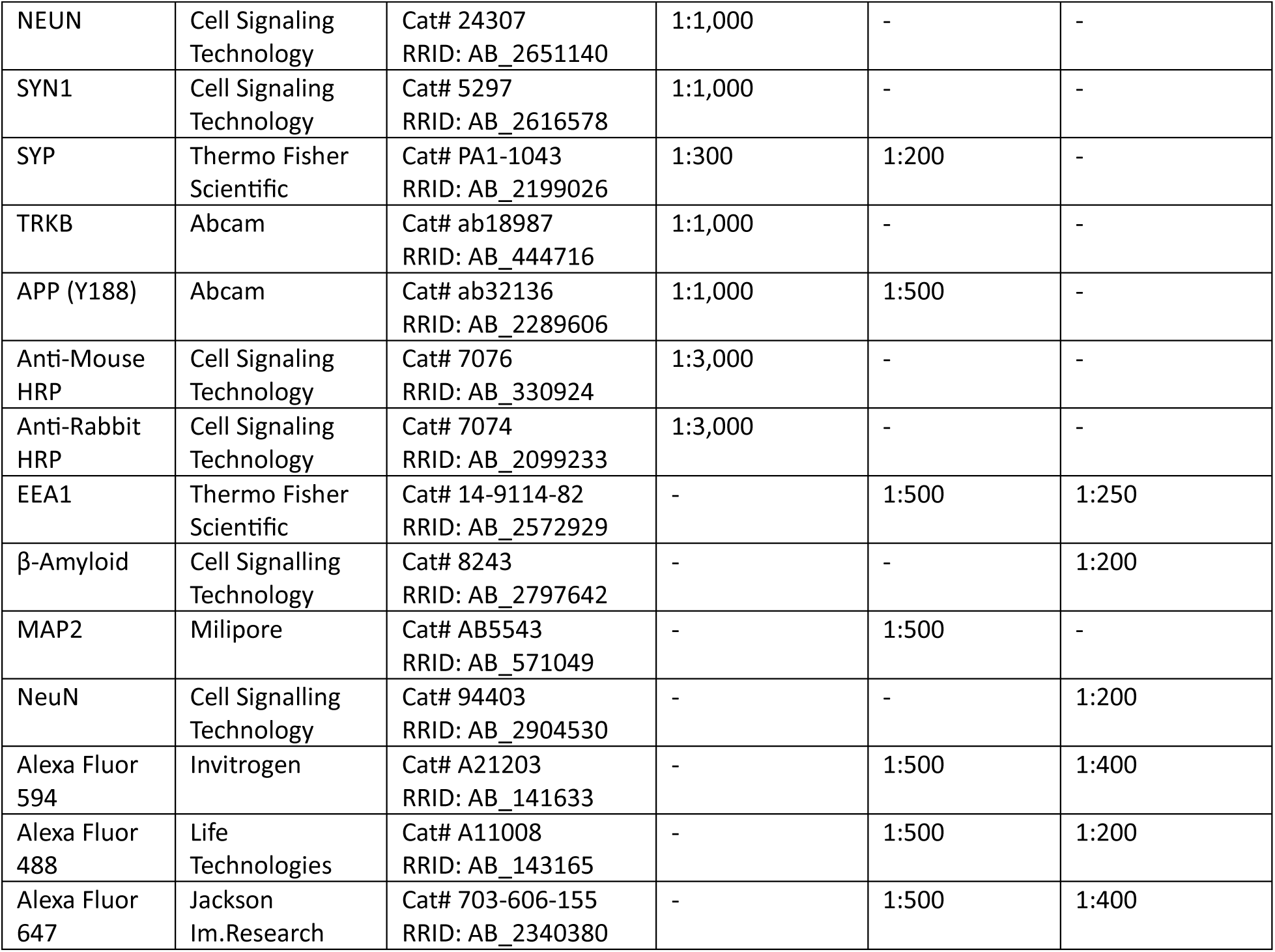
Media, primers and antibodies used in this study.

